# Cdc42 prevents precocious Rho1 activation during cytokinesis in a Pak1-dependent manner

**DOI:** 10.1101/2022.06.14.496145

**Authors:** Udo N. Onwubiko, Emma Koory, Sahara Pokharel, Hayden Roberts, Ahmad Mitoubsi, Maitreyi Das

## Abstract

Cytokinesis consists of a series of coordinated multi-step events that partition a dividing cell. Accurate regulation of cytokinesis is essential for proliferation and genome integrity. In fission yeast, these coordinated events ensure that the actomyosin ring and septum start ingressing only after chromosome segregation. How cytokinetic events are coordinated remains unclear. The GTPase Cdc42 is required for the delivery of certain cell wall-building enzymes while the GTPase Rho1 is required for activation of these enzymes. Here we show that Cdc42 prevents early Rho1 activation during cytokinesis. Using an active Rho-probe, we show that even though the Rho1 activators Rgf1 and Rgf3 localize to the division site in early anaphase, Rho1 is not activated until late anaphase, just before the onset of ring constriction. We find that loss of Cdc42 activation enables precocious Rho1 activation in early anaphase. Furthermore, this inhibition of Rho1 activation is dependent on the downstream Cdc42 effector Pak1 kinase. Disrupting pak1 function results in early Rho1 activation accompanied by precocious septum deposition and ring constriction. We provide functional and genetic evidence which indicates that Pak1 regulates Rho1 activation likely via the regulation of its GEF Rgf1. Our work proposes a mechanism of Rho1 regulation by active Cdc42 to coordinate timely septum formation and cytokinesis fidelity.

## Introduction

During cytokinesis, a mother cell is physically partitioned to generate two new daughter cells through a series of coordinated steps (Jordan and Canman, 2012; Pollard, 2010). The fission yeast model system divides via an actomyosin-based contractile ring, which is assembled in the medial region of the cell, as in animal cells (Balasubramanian et al., 2004; Pollard, 2010). The presence of a cell wall is essential for survival and cytokinesis in fungi. Thus, during cytokinesis in fission yeast a new cell wall known as the septum is synthesized to form the new ends of daughter cells. The septum primarily consists of linear β (1-3), branched β (1-6) glucan, and α-glucan components (Cortes et al., 2016; Munoz et al., 2013), and is formed in coordination with membrane ingression and ring constriction (Cortes et al., 2015; Cortes et al., 2016; Onwubiko et al., 2019). Failure to form a septum leads to cytokinetic failure since these cells cannot undergo ring constriction and membrane ingression (Arasada and Pollard, 2014; Cortes et al., 2015).

Cytokinesis begins with the assembly of protein nodes enriched in actin, formin Cdc12, type II myosin, F-BAR Cdc15, and IQGAP Rng2 at the division site (Padmanabhan et al., 2011; Wu et al., 2003). Under normal conditions, the ring is assembled as myosin heads interact with nucleated actin filaments to condense nodes into an actomyosin-based contractile ring (Wu et al., 2006). In fission yeast, ring constriction does not immediately follow ring assembly as in animal cells. Instead, once the ring assembles, it dwells or “matures” (Wu et al., 2003; Wu et al., 2006) while serving as a landmark for the recruitment of essential cytokinetic proteins such as Bgs1 (Arasada and Pollard, 2014; Vjestica et al., 2008; Wei et al., 2016). Membrane trafficking is required for the recruitment of septum synthesizing enzymes and furrow formation (Hercyk et al., 2019b; Vjestica et al., 2008; Wang et al., 2016). The septum consists of three layers made up of a primary septum sandwiched between two secondary septa (Pérez et al., 2018).

We have previously shown that Cdc42 promotes Bgs1 delivery and membrane trafficking at the division site (Onwubiko et al., 2021; Wei et al., 2016). Like all Rho-GTPases, Cdc42 is regulated by Guanine Nucleotide Exchange factors (GEFs) and GTPase Activating Proteins (GAPs), and is active when GTP-bound and inactive in the GDP-bound state. GEFs promote GTP binding, thus keeping the GTPase active while the GAPs increase GTP hydrolysis, thus promoting GTPase inactivation. In fission yeast, Cdc42 is sequentially activated by the GEFs Gef1 and Scd1 at the division site. Gef1 first activates Cdc42 during early anaphase, while Scd1 activates Cdc42 in late anaphase at the onset of ring constriction (Hercyk and Das, 2019b; Hercyk et al., 2019b; Wei et al., 2016). In the absence of *gef1* the initiation of septum formation and ring constriction is delayed (Wei et al., 2016).

In addition to Cdc42, other Rho GTPases also play a role in cytokinesis (Hercyk and Das, 2019a). The essential GTPase Rho1 is required for septum formation and cell wall integrity via the activation of cell wall building enzymes (Arellano et al., 1996). Septum synthesis and Rho1 activation occur once chromosomes segregation completes. The timing of septum synthesis is regulated by the Septation Initiation Network (SIN) pathway, a signal transduction cascade emanating from the microtubule-organizing center of the cell or the spindle pole body (SPB). The SIN pathway becomes active in early anaphase and is required for proper cytokinesis progression (Hou et al., 2000). The SIN is analogous to HIPPO signaling in mammalian cells, but the role of HIPPO in mammalian cytokinesis is yet to be clarified (Johnson et al., 2012). Genetic evidence indicates that the SIN pathway promotes Rho1 activation, which allows septum synthesis (Alcaide-Gavilán et al., 2014; Hou et al., 2000). Thus, the SIN pathway ensures septum synthesis is activated once chromosome segregation is successful.

Improper Cdc42 and Rho1 regulation have been shown to result in cytokinetic defects (Onishi et al., 2013; Onwubiko et al., 2021). It is unclear how Cdc42 and Rho1 coordinate septation during cytokinesis. Previous work in budding yeast suggested antagonistic roles for Rho1 and Cdc42, where Cdc42 inactivation and Rho1 activation are required for the completion of cytokinesis (Onishi et al., 2013). While Cdc42 has been shown to promote the delivery of Bgs1, Rho1 is the regulatory subunit required for the activation of Bgs1 (Arellano et al., 1996; Cabib et al., 1998). In larger eukaryotes, the role of Cdc42 activation has been reported mostly in meiotic division events such as polar body extrusion in oocytes, but not much is known about its role in cytokinesis in somatic cell division (Drechsel et al., 1997; Na and Zernicka-Goetz, 2006).

Here we investigate how Cdc42 and Rho1 regulate cytokinetic events. With the respective molecular probes designed to bind active Cdc42 and Rho1, we thoroughly compared the activity of these GTPases during cytokinesis in fission yeast. We report that Cdc42 activation precedes Rho1 activation at the division site. While Cdc42 activation occurs in early anaphase, Rho1 is activated in late anaphase, immediately before ring constriction is initiated. We show that Gef1-dependent activation of Cdc42 is required for inhibiting Rho1 activation in early anaphase. Further, we find that this inhibition is dependent on the Cdc42 effector Pak1 since, in *pak1* mutants, we observed abnormally early Rho1 activation and early septum deposits at the ring. Correspondingly, *pak1* overexpression rescues early activation of Rho1 at the division site in *gef1* mutants. To identify how Cdc42 prevents Rho1 activation we analyzed the functional relationship between *gef1* and various Rho1 regulators. The Rho1 GEFs localize to the division site in early cytokinesis even as Rho1 activation occurs in late cytokinesis indicating that their activity is regulated in early cytokinesis. We find that loss of the Rho1 GEF *rgf1* but not the essential GEF, *rgf3*, rescues premature Rho1 activation in both *gef1* and *pak1* mutants. Based on our data, we posit that premature Rho1 activation is prevented by Cdc42-dependent regulation of Rgf1. Together these results indicate that Cdc42 prevents Rgf1-driven Rho1 activation during early anaphase in a Pak1-dependent manner.

## Results

### Rho-probe detects Rho1 activation at cell tips and the division site

To study Rho1 activation, we designed a Rho-probe using the Rho-binding domain (RBD) of the protein kinase C, Pck2 modified from the RBD previously used in budding yeast (Kono et al., 2012; Yoshida et al., 2006). Briefly, we placed the promoter region of fission yeast *pck2* upstream of the *PCK2* RBD and a C-terminal fluorescent reporter and integrated the construct into the genome (Fig. 1Ai). Successful integration was confirmed by visualizing the localized Rho-probe signal in transformed wild-type cells, noting that these cells showed normal morphology with no aberrant phenotypes (Fig. 1Aii). The Rho-probe localized to the growing tips of interphase cells, and the division site of dividing cells (Fig. 1Aii and B, upper panel). It was previously reported that the Pck2p-RBD interacts with Rho1 and Rho2 GTPases (Arellano et al., 1999). To verify that our Rho-probe reported active Rho1 in cells, we expressed the Rho-probe in *rho2Δ* cells (Fig. 1B). We were still able to visualize bright Rho-probe signal at the cell tips, and at the division site in *rho2Δ* cells (Fig.1B). Thus, the Rho-probe reports active Rho1 localization in cells (Fig.1A, B).

**Figure 1.**
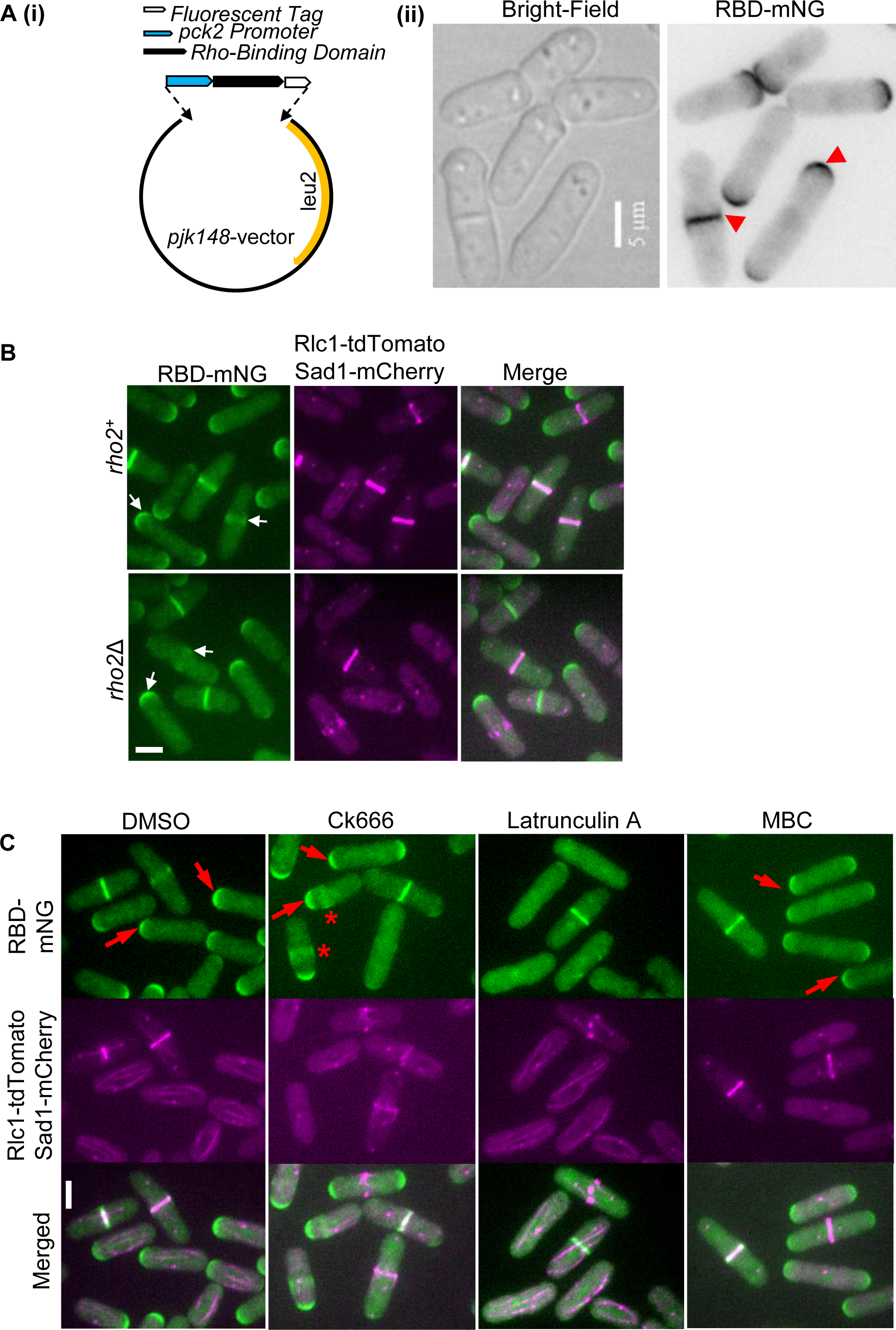
The Rho-probe detects Rho1 activation at cell tips and the division site. **A (i).** An Illustration of the Rho-probe design. **A (ii).** Rho-probe detects Rho activation in wild-type cells (red arrowheads) **B.** Rho-probe localization in the *rho2+* and *rho2Δ* strains, white arrows point to active Rho1 at the cell tips and division site. Rlc1-tdTomato marks the actomyosin ring and Sad1-mCherry marks the spindle pole bodies (SPB). **C.** Effect of cytoskeleton depolymerization on Rho probe localization in control YMD1045 strains, live cells were treated with DMSO, Ck666, Latrunculin A, and Methyl benzimidazol-2-yl-carbamate (MBC), (see methods for experimental details), Red arrows point to active Rho1 at cell tips, and asterisks mark ectopic Rho-probe localization, [Scale Bars, 5µm].

Since Rho1 activation occurs at the division site during cytokinesis, we asked if this required the actomyosin ring. The actomyosin ring can be disrupted in cells upon treatment with the drug latrunculin A (LatA). We find that in latA treated cells, Rho1 activation was not lost at the division site (Fig.1C, arrows). However, we did observe a loss of active Rho1 at the growing ends of interphase cells. Thus the actin ring does not promote Rho1 activation. Cytokinesis also requires endocytosis via branched actin networks. The drug CK666 has been shown to block the Arp2/3 complex and prevent branched actin assembly. We find that in cells treated with CK666 Rho1 activation was not lost at the division site. However, we observed ectopic Rho1 activation in interphase cells (Fig.1C, asterisks). We also depolymerized microtubules in cells treated with methyl benzimidazole-2yl carbamate (MBC). We did not observe any disruption in the localization pattern of active Rho1, however the signal appeared dampened at the cell tips and division site compared to DMSO-treated control cells (Fig. 1C). These results indicate that while at the division site the actin cytoskeleton is not required for Rho1 activation, it does play a role at the growth sites of interphase cells. Our data also indicates that Rho1 activation at the division site is differentially regulated than at the growing cell ends.

### Cdc42 is activated earlier while Rho1 is activated late in cytokinesis

We find that the Rho probe is only present at the division site of cells that also show septum deposition (Dashed white box, Fig. 2A). 3D-projections of the Rho-probe, the ring marker Rlc1-tdTomato, and the septum shows the spatial organization of the Rho-probe at the division site (Fig. 2A, Solid white boxes, top of the panel). The Rho-probe signal at the division site appears at the ring-membrane interface, overlapping with the calcofluor-stained septum (Fig.2A, 3D rings).

**Figure 2.**
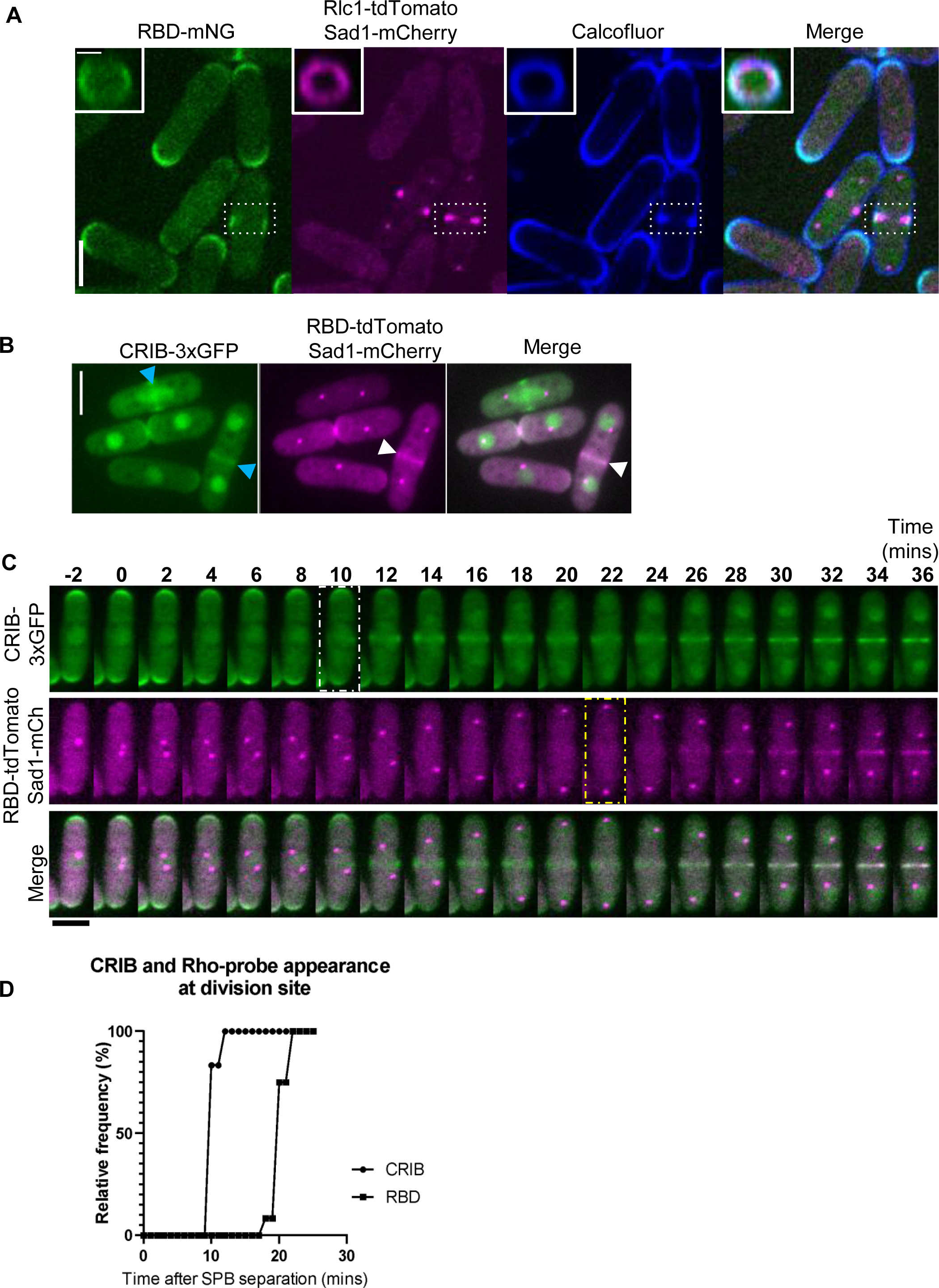
Cdc42 is activated earlier in cytokinesis while Rho1 is activated in late anaphase. **A.** Middle of Z-plane image shows active Rho, the ring (Rlc1-tdTomato) and septum (calcofluor white) at the division site in representative cells in cytokinesis (white box, Scale bar 5µm). 3D-Projections (Rings enclosed in white box, top panel) of division site (white box) shows concentric rings of active Rho (green), the actomyosin ring (Magenta), and the septum (blue), [Scale bar for 3D projections, 2µm]. **B.** Localization of active Cdc42 (CRIB-3xGFP, blue arrowheads), and active Rho-probe in cells (RBD-mNG, white arrowheads) [Scale Bars, 5µm]. **C.** Time-lapse series of a representative cell, time of Cdc42 activation (white box) and Rho activation (yellow box) during cytokinesis [Scale Bar for cells, 5µm] **D**. Quantification of the timing of Cdc42 and Rho activation at the division site during cytokinesis (n=12 cells), p<0.0001 determined by one way ANOVA followed by Tukey’s HSD test].

To investigate Rho1 and Cdc42 dynamics during cytokinesis, we performed live imaging of cells simultaneously expressing the active Cdc42 probe CRIB-3xGFP (Das et al., 2012; Tatebe et al., 2008; Wei et al., 2016) and the Rho-probe, RBD-tdTomato in wild-type cells (Fig. 2B). We observe that while some cells display RBD-tdTomato and CRIB-3xGFP simultaneously at the division site, others only display CRIB-3xGFP (Fig. 2B). Time-lapse imaging of cells simultaneously expressing CRIB-3xGFP and RBD-tdTomato during cytokinesis reveals that Cdc42 was activated at the division site ∼10minutes after the duplication of the spindle pole bodies (SPBs), while Rho1 is activated ∼20minutes after SPB duplication (Fig. 2B). Following the previously described timeline of cytokinetic events in fission yeast (Wu et al., 2003), we observe Cdc42 activation in early anaphase, at the time of ring assembly. In contrast, we find that the Rho-probe localizes to the division site in late anaphase, at the onset of ring constriction, suggesting that Rho1 is activated later in cytokinesis (Fig. 2 C, D, S1A). These observations are consistent with the fact that septum formation is activated by Rho1, and occurs in late anaphase.

Rho1 is activated by the GEFs Rgf1, Rgf2, and Rgf3 (Mutoh et al., 2005; Tajadura et al., 2004), all of which promote cell wall integrity (Fig. 3A). We asked if the delay in Rho1 activation was due to a delay in the localization of its GEFs at the division site. Previously, it was reported that loss of *rgf1*or *rgf2* does not disrupt cell viability, but *rgf1Δ* cells display severe cell wall defects, and *rgf1Δ rgf2Δ* is lethal(Mutoh et al., 2005). Rgf3 is the primary cytokinetic GEF for Rho1, and loss of *rgf3* is inviable (Tajadura et al., 2004). We imaged live cells expressing fluorescently labeled Rgf3 and Rgf1. Both Rgf1-GFP and Rgf3-eGFP localize to the division site in early anaphase (Fig. 3B, C) at the time when active Rho1 is absent from the division site. While we could not acquire time-lapse series to assess the exact timing of Rho1 GEF localization, we quantified Rgf1 and Rgf3 localization during cytokinesis by measuring the distance between the mitotic spindle pole bodies (SPBs). SPB duplication and movement from mitosis onset serve as a conventional timer for cytokinesis (Nabeshima et al., 1998; Pollard and Wu, 2010).

**Figure 3.**
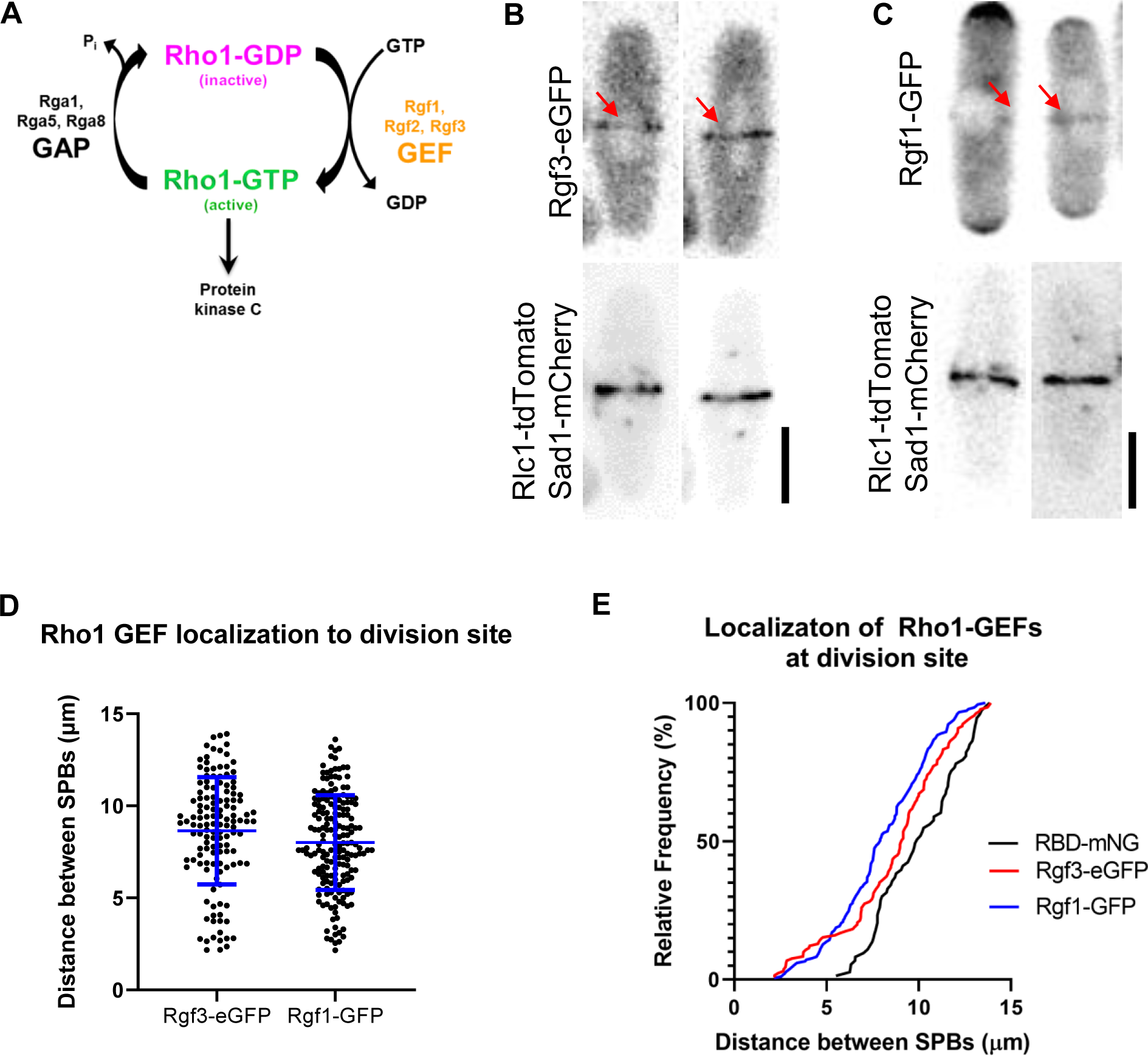
Rho1-GEFs Rgf1 and Rgf3 localize to the division site early during cytokinesis. **A.** A schematic of the Rho1-GTPase activation cycle showing the known GEFs and GAPs. **B.** Localization of Rgf3-mEGFP and **C.** Rgf1-GFP to the division site (Red arrows) [Scale Bar, 5μm]. **D.** Quantification of the distance between the spindle pole bodies (SPBs) for strains as indicated, [n≥94 cells for each genotype quantified; Error bars represent the standard deviation]. **E.** Frequency plot of the shortest SPB distances at which Rgf3 and Rgf1 are present at the division site. For these quantifications, shorter SPB distances represent an earlier time-point during cytokinesis. [n≥90 cells for each genotype quantified; Error bars represent the standard deviation].

Cytokinesis progression is therefore proportional to the distance between mitotic spindles up until late anaphase when ring constriction and septum formation begin. Here we labeled the SPBs with Sad1m-Cherry in cells expressing the fluorescently tagged GEFs and computed SPB distances at which the GEF localizes to the division site (Fig. 3B, C, D). In an asynchronous population of cells, we find that Rgf1 and Rgf3 localize as early as the time of ring assembly at an average SPB distance of 4-5μm (Fig. 3D). A comparison of the cumulative frequency distribution of the localization of the two GEFs with increasing SPB distance suggested that the GEFs localize to the division site simultaneously (Fig. 3E). However, while the GEFs localize early to the division site, the Rho probe was visible only at a greater SPB distance suggesting a time delay between GEF localization and Rho1 activation. Thus, while Rho1 GEFs Rgf1 and Rgf3 localize to the division site in early anaphase, they are unable to activate Rho1 at this stage.

### Active Cdc42 inhibits Rho1 activation in early anaphase

Next, we asked what prevents early Rho1 activation. Since Cdc42 is activated before Rho1, we asked if active Cdc42 prevents early Rho1 activation. As previously shown, Cdc42 activation in early anaphase is dependent upon its GEF Gef1 (Wei et al., 2016). To test if active Cdc42 prevents Rho1 activation during cytokinesis we looked at Rho1 activation in *gef1Δ* cells. Time-lapse imaging of *gef1+* and *gef1Δ* cells expressing the Rho-probe reveal that *gef1Δ* causes premature Rho activation in early anaphase (Fig. 4A). We find that ∼100% of *gef1Δ* cells display early Rho-probe localization at the division site at ∼12 minutes after SPB duplication compared to ∼20 minutes in *gef1+* control cells (Fig. 4B). Furthermore, we find that Rho-probe normally localizes to the division site at SPB distances of greater than ∼7µm in *gef1+* control cells, while in *gef1Δ* cells they appear early at an SPB distance of ∼4 µm or less (Fig. 4C).

**Figure 4.**
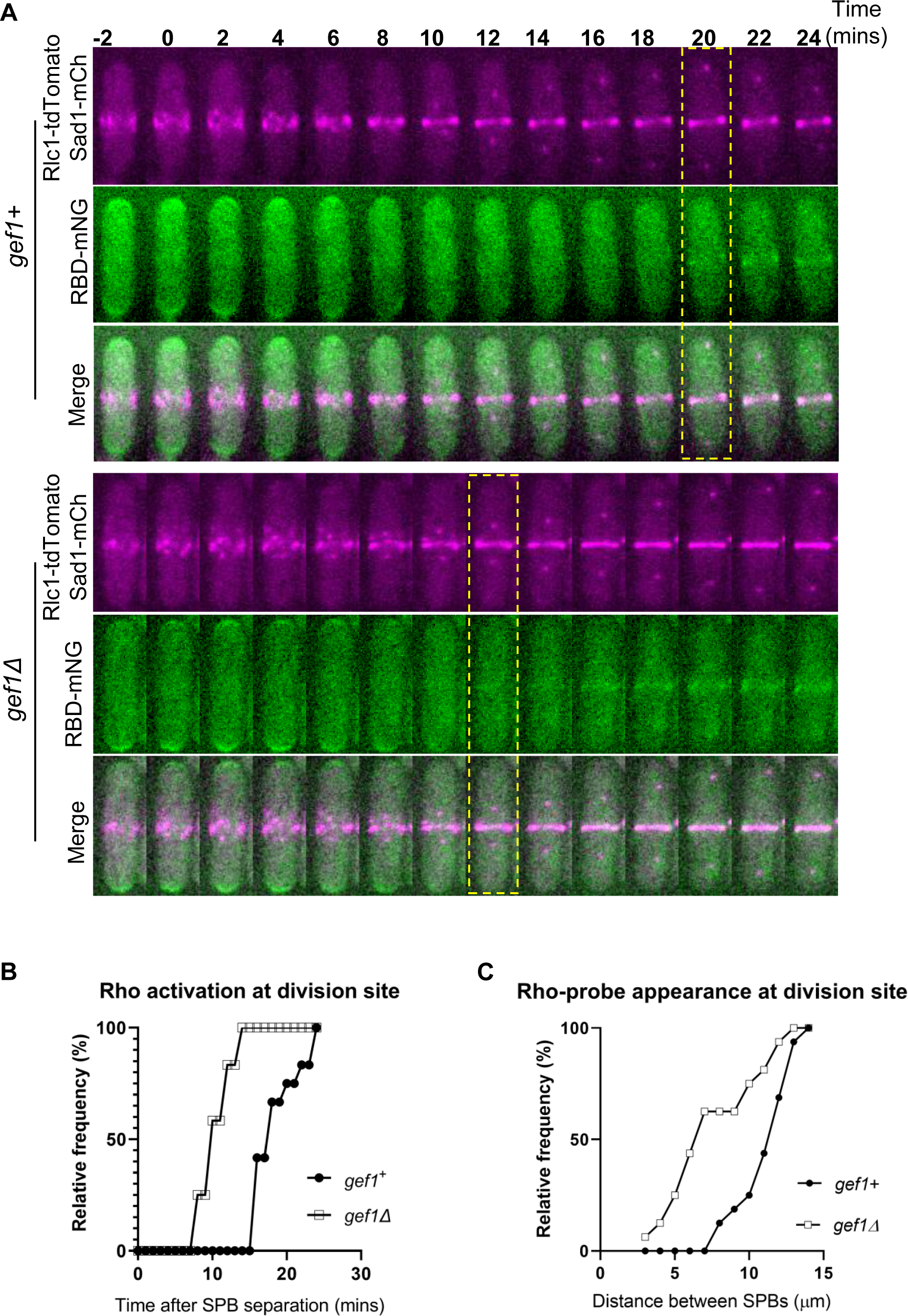
Loss of *gef1* results in early Rho1 activation early in cytokinesis. **A.** Time-lapse montage shows the time of Rho activation at the division site in representative *gef1+* and *gef1Δ* cells (yellow box), Time=0 marks the time of SPB separation, and onset of cytokinetic events, [the statistical significance for these movies are shown in Figure 9C, Scale Bar, 5µm]. **B.** Outcome plot shows frequency of Rho-activation over time during cytokinesis from time-lapse movies of strains as indicated [n= 12 cells from time-lapse movies]. **C.** Outcome plot shows frequency distribution of SPB distances at which Rho-activation at the division site is observed in acquired still images of *gef1+* and *gef1Δ* cells. [n=16cells per indicated strains].

To test whether the Gef1-dependent inhibition of Rho1 activation in early cytokinesis was mediated by Cdc42 activation we assessed the localization of the Rho-probe in cells expressing the constitutively active *cdc42G12V* allele. Previously, we have shown that cytokinetic defects in *gef1* mutants can be rescued by moderate expression of constitutively active *cdc42G12V* (Wei et al., 2016). With the medium-strength thiamine-repressible promoter, *nmt41*, we moderately expressed *cdc42G12V* in *gef1+* and *gef1Δ* cells containing the Rho-probe. Experimental controls were *gef1+* and *gef1Δ* cells expressing the empty pjk148 vector with the Rho-probe. All cells expressed the ring marker Rlc1-GFP, and SPB marked by Sad1m-Cherry. As performed in Figures 3E, and 4C, the distance between the SPBs was used to identify the stages of cytokinesis at which the Rho-probe was detected at the division site. We find that the Rho-probe localizes to the division site in late anaphase in *gef1+* pjk148 empty control cells and is early in *gef1Δ* pjk148 empty cells (Fig. 5A, B). However, the expression of *cdc42G12V* in *gef1Δ* cells reverts Rho-activation at the division site to late anaphase, similar to *gef1+* controls (Fig.5A, B). We quantified the percentage of non-constricting actomyosin rings with Rho-probe localization and find a significant increase in *gef1Δ* cells compared to *gef1+* controls, and this is rescued by *cdc42G12V* expression in *gef1Δ* (Fig. S1D). We also observed that the Rho-probe signal is diminished in *cdc42G12V* expressing cells at the cell tips and division site (Fig. 5A). We quantified the mean fluorescence intensity of the Rho-probe signal at the division and confirm that it is indeed significantly reduced in *cdc42G12V* expressing cells (Fig. 5C).

**Figure 5.**
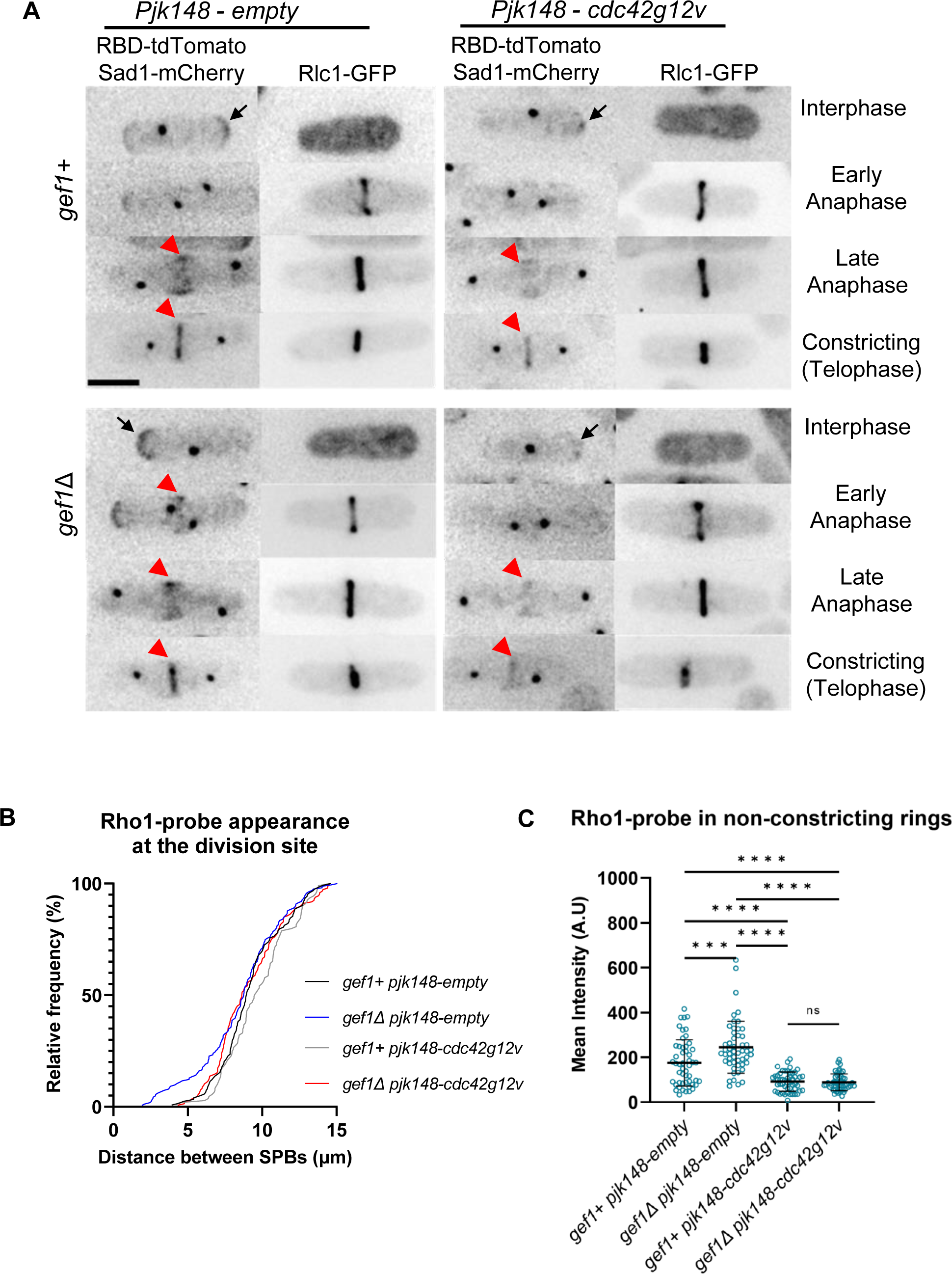
Constitutively active *cdc42* mutants rescue early Rho1 activation in *gef1Δ* mutants. **A.** Rho1 activation at the division site (red arrowheads) and cell tips (black arrows) of representative *gef1^+^* and *gef1Δ* cells, transformed with the empty vector pJK148, or expressing constitutively active *cdc42G12V* [Scale Bar, 5µm]. **B.** Outcome plot shows the frequency distribution of SPB distances for which active Rho1 is observed at the division site in the indicated strains, [n=160 cells per strain indicated]. **C.** Quantification of the mean fluorescence intensity in strains [n≥60 cells per strain; Statistical significance determined with 2-way ANOVA, with Tukey’s multiple comparisons post hoc test, ***p≤0.0002 ****p≤0.0001; n.s - not statistically significant; Error bars represent standard deviation].

We questioned whether early Rho-probe localization in *gef1Δ* cells during cytokinesis was due to Rho1 or Rho2 activity. To verify this, we assessed the localization of the Rho-probe in *rho2Δ* cells (Fig. S1B). We find that while Rho-probe localizes in late anaphase in *gef1+ rho2+* control cells, it still localizes early in *gef1Δ rho2Δ* double mutants (Fig. S1B, C.). Hence, early Rho-probe localization in *gef1Δ* cells is indeed due to early Rho1 activation. Collectively, these data indicate that active Cdc42 prevents premature Rho1 activation in early anaphase. As per our observations, we refer to the Rho-probe signal at the division site as active Rho1 from here onwards.

Genetic evidence indicates that Rho1 activation requires activation of the SIN pathway. We asked whether early Rho1 activation in *gef1Δ* mutants was sufficient to bypass the requirement for SIN-dependent activation. Using the Rho-probe, we assessed Rho1 activation in a temperature-sensitive *sid2* mutant (*sid2-250*) which disrupts SIN function at the restrictive temperature (36°C). As previously shown (Feoktistova et al., 2012), SIN inactivation resulted in elongated, non-dividing cells (Fig. S2A.). We find that under permissive conditions (25°C) Rho1-probe localizes to the division site in both *gef1+ sid2-250* and *gef1Δ sid2-250* cells (Fig. S2A, B). However, SIN inactivation at 36°C abolished Rho-probe localization from the division site in *gef1+ sid2-250* and *gef1Δ sid2-250* cells (Fig. S2A, B) suggesting that the SIN is required for Rho1 activation during cytokinesis. Interestingly, these mutants were still able to activate Rho1 at the cell tips suggesting that the SIN pathway is only required for Rho1 activation at the division site.

### Disruption of Pak1 kinase function results in early Rho1 activation during cytokinesis

The p21-activated kinase (Pak1) is a known downstream effector of Cdc42 in fission yeast (Magliozzi et al., 2020; Ottilie et al., 1995). In animal cells, the RhoA GTPase GEFs are differentially regulated by PAK kinases (Alberts et al., 2005; Zenke et al., 2004). It has also been shown that Pak1 localizes to the division site in early cytokinesis (Magliozzi et al., 2020), via its interaction with active Cdc42. Given its early localization to the division site, we wondered if Cdc42-dependent inhibition of Rho1 activation was due to Pak1 function. We reasoned that if Pak1 was responsible for blocking Rho1 activation, then loss of *pak1* function should allow early Rho1 activation at the division site. To evaluate this idea, we assessed Rho1 activation in the temperature-sensitive *pak1* mutant (*orb2-34*) (Verde et al., 1995) which is labeled onwards as *pak1-ts* for clarity. We first quantified the appearance of the Rho-probe at the division site in the temperature-sensitive *pak1* mutants. At the permissive temperature, Rho1 activation at the division site was normal in late anaphase in *pak1+* cells (Fig.6A). In *pak1-ts* mutants at 25°C, Rho1 activation also occurred a bit earlier that in *pak1+* cells as determined by the distance between SPBs and the appearance of assembling rings (Fig. 6B). At 35.5°C however, the active Rho1-probe localized in early anaphase in most *pak1-ts* mutant cells (Fig. 6A, B). To assess the timing of Rho1 activation during cytokinesis, we measured the distance between the Sad1-mCherry-labeled SPBs in *pak1+* and *pak1-ts* cells (Fig. 6B). At 25°C, *pak1+* and *pak1-ts* cells show Rho1 activation late in anaphase as determined by longer SPB distances (Fig. 6A, B). In contrast, at 35.5°C *pak1-ts* mutants displayed early Rho1 activation at the division site as determined by the short SPB distances. Thus, disruption of *pak1* function enables early Rho1 activation in cytokinesis. While we observed RBD at a shorter SPB distance in *pak1* mutants under restrictive conditions, we also note that these mutants are smaller in size. However, the small cell size does not impact the SBB distance at which the Rho-probe appears at the division site. This is because even under permissive conditions, *pak1* mutants are smaller in size, and in these cells, the Rho-probe does not appear at the division site at a relatively short SPB distance (Fig. 6B).

**Figure 6.**
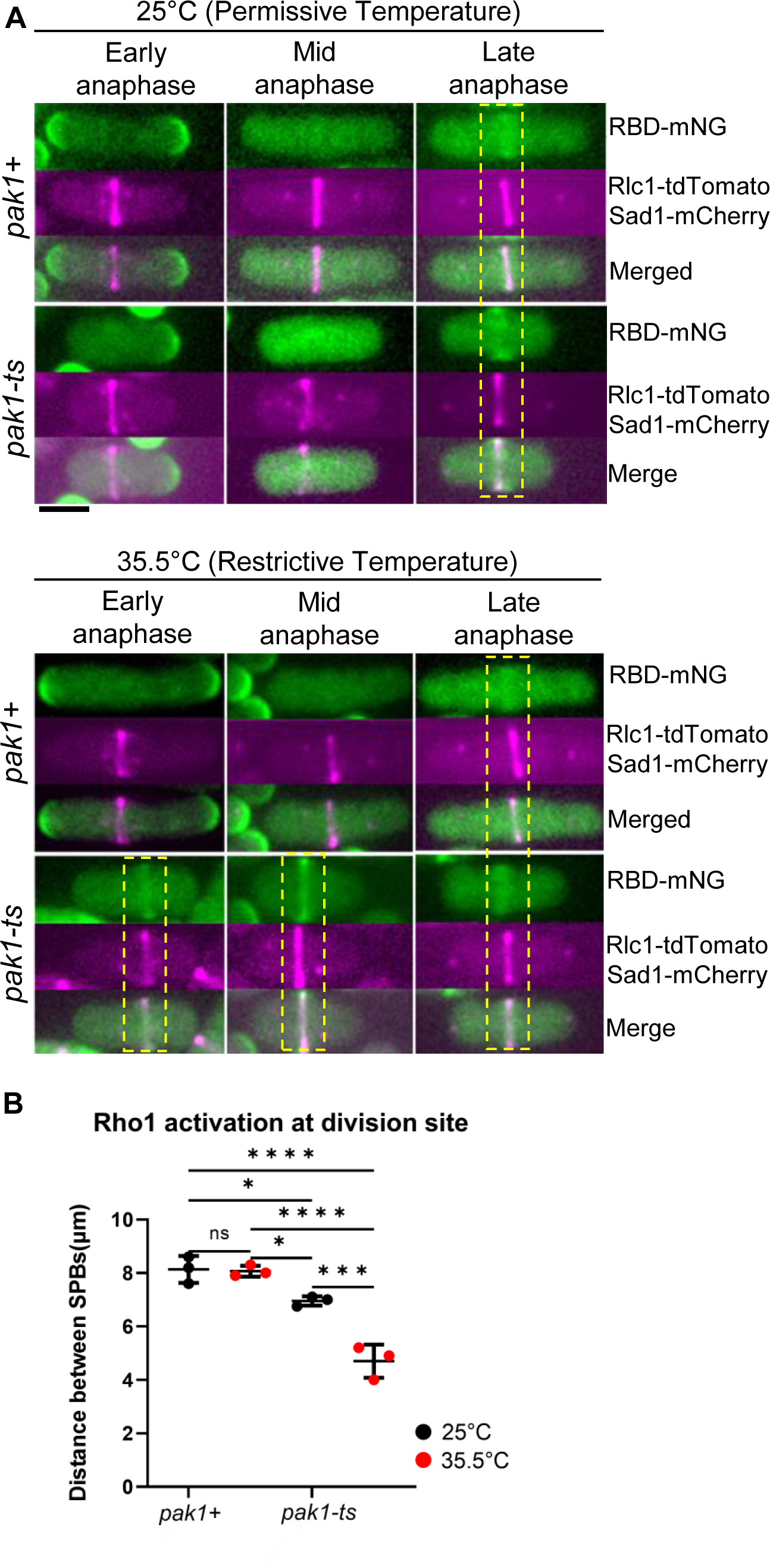
Hypomorphic *pak1* mutant displays early Rho1 activation in cells during cytokinesis. **A.** Rho1 activation in *pak1+* (*orb2+*) and *pak1-ts* (*orb2-34*) strains at the permissive temperature (25°C), and restrictive temperature (35.5°C). Yellow boxes highlight the stage of cytokinesis in which Rho1 activation is observed at the division site in the indicated strains, [Scale Bar 5μm]. **B.** Outcome plot shows frequency distribution of the quantified distance between the SPBs at which Rho1 activation at the division site is observed in all conditions shown [N= 3 representative experiments. Data points on the graph represent the first quartile of SPBs distances measured in the indicated strains; Statistical significance between strains is determined by one-way ANOVA followed by Tukey’s HSD test, *p≤0.03, ***p≤0.0008, ****p≤0.0001; n.s - not statistically significant; Error bars represent standard deviation].

Next, to further confirm that inhibition of Rho1 is indeed dependent on Pak1 kinase, we tested whether *pak1* overexpression disrupts Rho1 activation in dividing cells even in the absence of *gef1*. Utilizing the high-strength thiamine repressible promoter nmt1, (Javerzat et al., 1996), we assessed Rho1 activation during cytokinesis. The nmt1-3xHA-*pak1* allele (MBY3451) was either expressed or repressed in *gef1+* and *gef1Δ* cells expressing the Rho-probe, Sad1m-Cherry, and Rlc1-tdTomato (Fig. 7A). Experiments were performed by growing cells in the presence - (*pak1-*), and absence (*pak1OE*) of thiamine. The high-strength *nmt1* promoter does show leaky expression even in the presence of thiamine (Javerzat et al., 1996; Wei et al., 2016). Thus *gef1+ pak1-* cells behave similar to wild type and show Rho1 activation in late anaphase, (Fig. 7B, C). However, early Rho1 activation at the division site observed in *gef1Δ* cells was rescued upon *pak1OE*, restoring it to late anaphase (Fig. 7A, B). We also observed an overall decrease in mean intensity of the Rho-probe in *pak1OE* cells compared to *pak1-* cells (Fig. 7A, C), in agreement that overexpression of *pak1* blocks Rho1 activation. Together, these data suggest that Gef1 mediates Rho1 inhibition in early anaphase via the Pak1 kinase.

**Figure 7.**
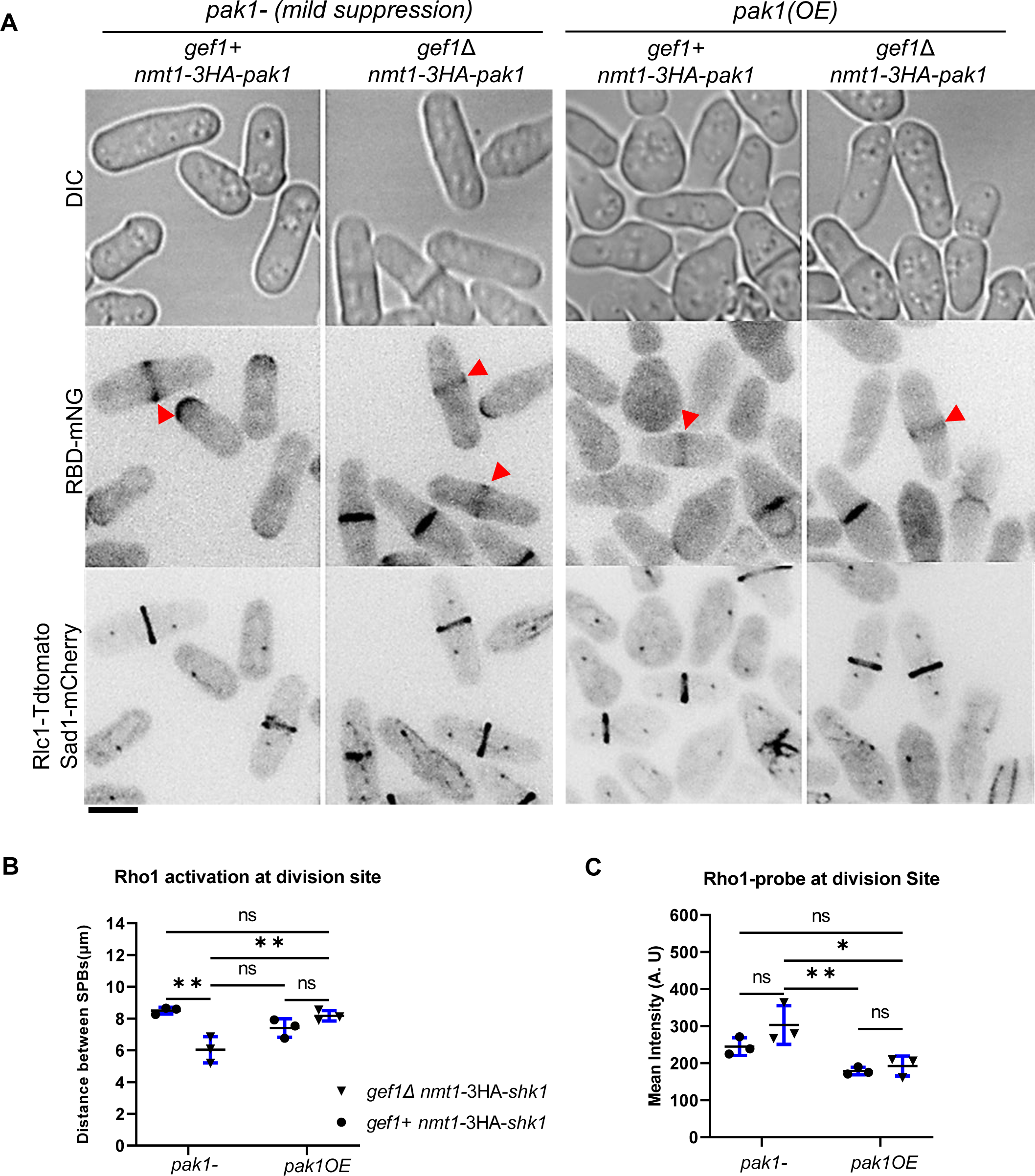
Overexpression of *pak1* (*pak1*OE) rescues early Rho1 activation in *gef1Δ* cells. **A.** Rho1 activation during cytokinesis in *gef1+* and *gef1Δ* cells expressing nmt1-3HA-*pak1*, in thiamine repressing (*pak1-*), or overexpressing (*pak1*OE) conditions (see methods). Red arrowheads point to division sites displaying the active Rho1-probe [Scale Bar 5μm]. **B.** Quantification of the cells with active Rho1-probe at the division site during cytokinesis progression as indicated by the SPB distance. Data points on graph represent first quartile of measurements obtained from N= 3 replicate experiments. [Statistical significance between strains determined by one-way ANOVA followed by Tukey’s HSD test, **p≤0.005; n.s - not statistically significant; Error bars represent standard deviation]. **C.** Quantification of the mean fluorescence intensity of the active Rho1-probe localized to the division site of mentioned strains and conditions. [N= 3 replicate experiments; Statistical significance between strains determined by one-way ANOVA followed by Tukey’s HSD test, *p≤0.01, **p≤ 0.006; n.s- not statistically significant; Error bars represent standard deviation].

### Loss of *gef1* rescues *rgf3* repression-induced lethality

Based on our observation that Rgf1 and Rgf3 localize to the division site in early anaphase, even before the ring is fully assembled (Fig.3B, C), we wondered whether the localization of either of these Rho-GEFs was either enhanced or early in *gef1* mutants. However, in comparing Rgf3-eGFP and Rgf1-GFP localization in *gef1+* and *gef1Δ* cells using the distance between SPBs as the marker for cytokinesis progression, we saw that both GEFs localize in early cytokinesis in *gef1+* and *gef1Δ* cells (Fig. S3A, B). The mean intensity of Rgf1 and Rgf3 at the division site in *gef1+* and *gef1Δ* cells also remained similar (Fig. S3C, D). Thus neither the localization nor intensity of either Rgf3 or Rgf1 is changed in the absence of *gef1*. This result suggests that regulation of Rho1-GEF activity prevents Rho1 activation in the early anaphase.

Given that Rgf3 is the only Rho1 GEF to localize specifically to the division site but not the cell tips, (Morrell-Falvey et al., 2005; Mutoh et al., 2005), we asked if active Cdc42 inhibits Rgf3-dependent Rho1 activation in early cytokinesis. Loss of *rgf3* results in cell lysis and death (Tajadura et al., 2004). To test if Rgf3 was responsible for early Rho1 activation, we repressed *rgf3* expression via the low strength thiamine-repressible promoter, *nmt81*, in *gef1+* and *gef1Δ* cells. We first assessed the viability of *rgf3*-depleted (*rgf3-*) cells in the absence of *gef1* by performing a growth assay of *gef1+ rgf3+*, and *gef1Δ rgf3-* cells. Cells were subjected to *rgf3*-expressing or thiamine-repressing conditions in growth media at 25°C. As expected, *gef1+ rgf3-* cells showed lethality under *rgf3*-repressing conditions, however, *gef1Δ rgf3-* survived under these conditions (Fig. 8A). To corroborate our observations, we analyzed Rho1 activity at the division site in *gef1Δ rgf3-* cells. Repression of *rgf3* (*rgf3-*) resulted in lysed and dying cells as shown previously (Tajadura et al., 2004). In *gef1+* cells, repression of *rgf3* resulted in lysis of ∼58% of cells in an asynchronous population (Fig. 8B, black arrowheads, C). Loss of *gef1* rescued the lethality of *rgf3*-cells, thus corroborating our growth assay results (Fig. 8B, C). Moreover, *rgf3-depletion* did not disrupt Rho1 activation at the onset of ring constriction in *gef1+* and *gef1Δ* cells (Fig. 8B, D). This indicates that the loss of *gef1* rescues the lethality of *rgf3-* cells, and early Rho1 activation in *gef1* mutants is not Rgf3-dependent. This suggests that Gef1-mediated repression of Rho1 activity occurs via another mechanism. Thus, Rgf3 may function to promote septation integrity at later stages in cytokinesis but is not responsible for early Rho1 activation in cytokinesis.

**Figure 8.**
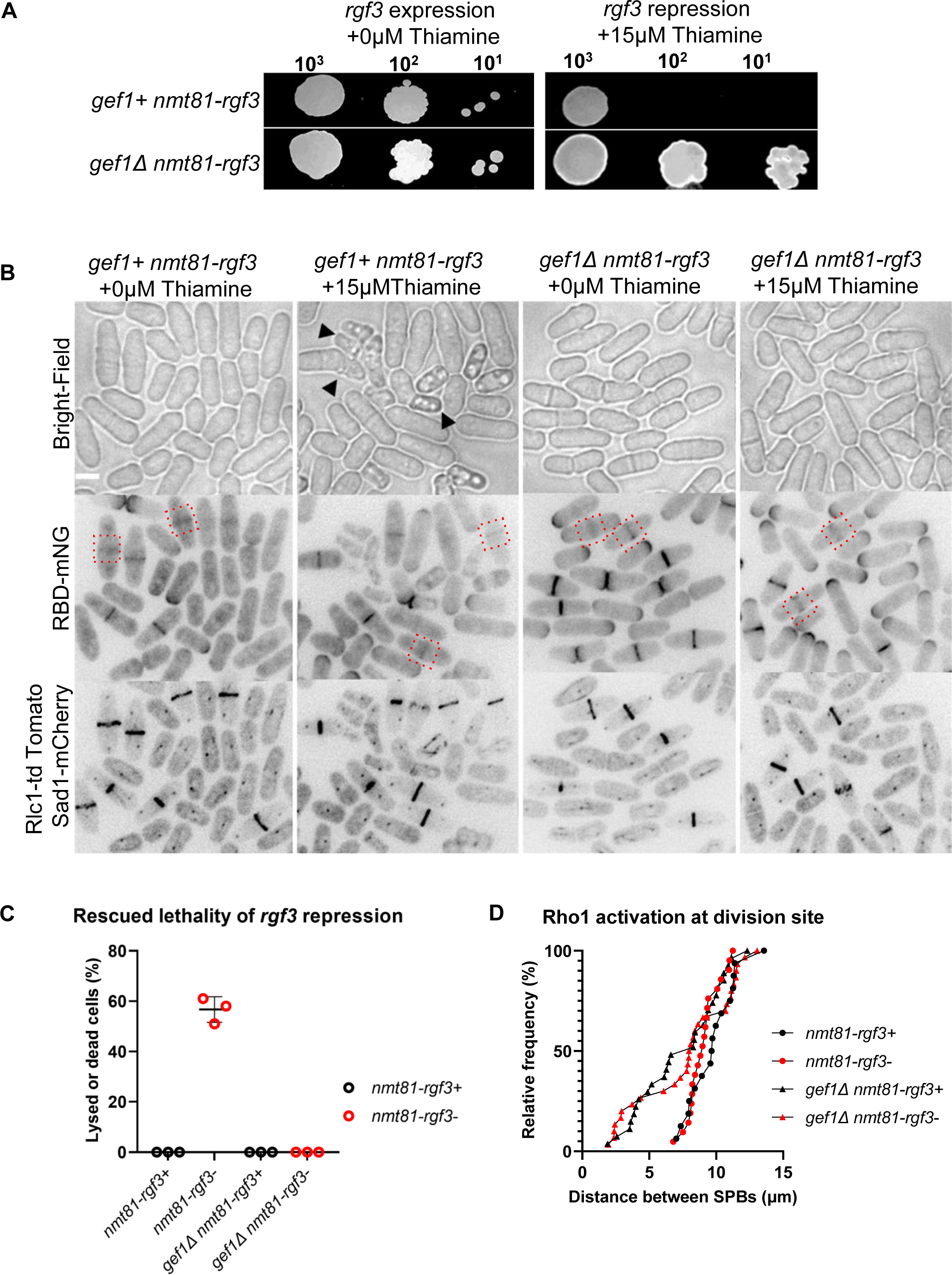
Loss of *gef1* rescues lethality of *rgf3*-repressed strains. **A**. Rescue of *rgf3-*repression lethality in *gef1Δ* cells as shown by a spot assay on supplemented EMM media with or without 15µM thiamine for the strains as indicated. **B.** Rho1 activation at the division site (red dashed boxes) under *rgf3-*repressed and expressed conditions in the indicated strains. Black arrows point to dying or lysed *rgf3-*depleted cells [Scale Bar, 5µm]. **C.** Quantification of cell lysis in the indicated strains [For all quantifications, N=3 experiments; Error bars represent standard deviation] **D.** Outcome plot shows the distance between SPBs at which Rho1 activation is observed at the division site of the indicated strains and conditions. [For all quantifications, n=30cells per indicated strains; Error bars represent standard deviation].

### Loss of *rgf1* rescues premature Rho1 activation in *gef1* mutants

While Rho1 is activated by the GEFs Rgf1/2/3, it is inactivated by the GAPs Rga1/5/8 (Calonge et al., 2003; Nakano et al., 2001). Of these Rho1 GAPs, Rga5 has been implicated in proper Rho1 function during cytokinesis (Calonge et al., 2003). Loss of *rga5* function has been reported to rescue septation defects in cells where Rho1 activity is impaired and is lethal in cells where Rho1 is overexpressed (Alcaide-Gavilán et al., 2014; Calonge et al., 2003). To address if Cdc42 inhibited Rho1 activation by promoting Rga5 GAP function, we compared Rho1 activation at the division site in *gef1+*, *rga5Δ*, *gef1Δ,* and *gef1Δ rga5Δ* cells. We expect that if Rga5 is responsible for Cdc42-dependent Rho1 inhibition in early cytokinesis, then *rga5Δ* should phenocopy *gef1Δ*. We find that in *rga5Δ* Rho1 activation occurs earlier than in *gef1+* cells (Fig. 9C), but not as early as in *gef1Δ* cells. Thus Rga5 alone cannot account for the premature Rho1 activation in *gef1Δ* cells. This suggests that Gef1 prevents Rho1 activation in early anaphase via other Rho1 regulators.

**Figure 9.**
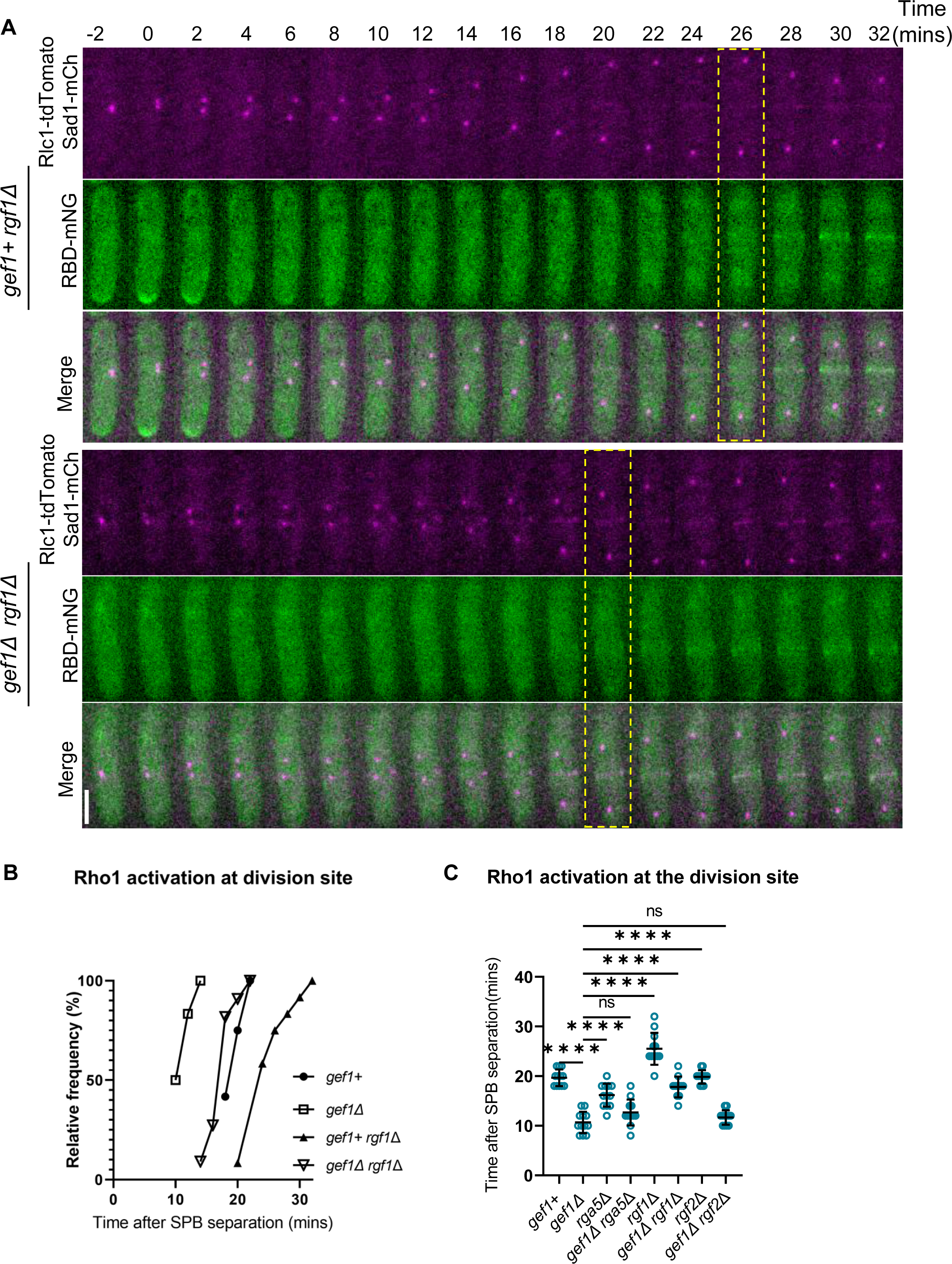
Loss of *rgf1* rescues premature Rho1 activation in *gef1* mutants. **A.** Time-lapse of representative *gef1+ rgf1Δ* and *gef1Δ rgf1Δ* cells shows the time of Rho1 activation at the division site during cytokinesis (yellow box) [Scale Bar 5μm. The controls *gef1+ rgf1+* and *gef1Δ rgf1+* are shown in Supplemental Figure 4 (Fig.S4). **B.** Outcome plot shows the frequency of the time at which Rho1 activation occurs at the division site during cytokinesis as determined from time-lapse movies as shown in A. Time=0 marks the onset of cytokinetic events, [n=12 cells per strain]. **C.** Quantification of the timing of Rho1 activation during cytokinesis from movies of the strains indicated, [n=12 cells per strain; Statistical significance between strains determined by one-way ANOVA followed by Tukey’s HSD test ****p≤0.0001; n.s *-* not statistically significant; Error bars represent standard deviation].

Rho1 GEFs differ in their spatial dynamics during ring constriction. Unlike Rgf3 which localizes at, and constricts with the actomyosin ring, Rgf1 and Rgf2 ingress with the membrane furrow and septum (Morrell-Falvey et al., 2005; Mutoh et al., 2005). Active Rho1 at the division site appears diffuse and condenses into a band-like appearance as constriction begins (Fig. 2A, C). Rho1 may therefore be differentially activated by its GEFs at the division site. Our observation that Rho1 activation remained early in *gef1Δ* cells depleted of *rgf3* (Fig. 6B, D) suggests that another GEF is responsible for this activation in early cytokinesis. Therefore, we tested whether the other Rho1 GEFs *rgf1* and *rgf2* were required for early Rho1 activation in *gef1Δ* cells. While *rgf1Δ* is viable, these cells display severe cell wall and morphology defects (Tajadura et al., 2004). On the other hand, *rgf2Δ* cells do not show any obvious defect, but the *rgf1Δ rgf2Δ* double mutant is not viable (Morrell-Falvey et al., 2005; Mutoh et al., 2005). This indicates that Rgf1 and Rgf2 are redundant with Rgf1 being the primary Rho1 GEF. We asked whether early Rho1 activation in *gef1Δ* cells was Rgf1-dependent. We assessed the Rho-probe in *gef1+*, *gef1Δ, rgf1Δ*, and *gef1Δ rgf1Δ* cells with time-lapse confocal microscopy. The Rho-probe localized normally at the onset of ring constriction in *gef1+ rgf1+* control cells (Fig. 9A). As expected, Rho1 activation also occurred early in *gef1Δ* (Fig. 9B and Fig. 4). Rho1 activation at the division site was delayed in *rgf1Δ* cells compared to the *gef1+* controls (Fig. 9A, B).

Interestingly, loss of *rgf1* prevents premature Rho1 activation in *gef1Δ* cells restoring it to normal in late anaphase (Fig.9A, B). This suggests that Rgf1 is responsible for early Rho1 activation in *gef1Δ* cells. We observed that Rho1 activation in *rgf1Δ* cells was delayed much more than in *rgf1Δ gef1Δ* cells (Fig. 9A-C) indicating that in the *gef1Δ* in addition to Rgf1, another regulator is responsible for early Rho1 activation. We asked if this regulator could be the other GEF Rgf2 which has been shown to behave similarly to Rgf1 (Morrell-Falvey et al., 2005). We find that *rgf2Δ* did not alter Rho1 activation at the division site (Fig. S4A, 9C), likely due to the presence of Rgf1. Accordingly, we see that *gef1Δ rgf2Δ* double mutants show early Rho1 activation at the division site similar to *gef1Δ* cells (Fig. S4A, B,C). While we were unable to assess the Rho-probe in the *rgf1Δ rgf2Δ* double mutants due to its lethality, our observations suggest that apart from Rgf1 early Rho1 activation in *gef1Δ* cells is either due to activation of Rgf2 or due to inhibition of Rga5.

Given that Rho1 activation at the division site occurred early in *pak1-ts* cells at the restrictive temperature, we posit that Pak1 prevents Rgf1-dependent Rho1 activation in early anaphase. We tested our idea by analyzing Rho1 activity in *rgf1Δ pak1-ts* double mutant. If indeed Pak1 regulates Rgf1, then the loss of *rgf1* in *pak1-ts* mutants should rescue early Rho1 activation at the division site. As performed earlier, we imaged the *rgf1+*, *pak1-ts*, *rgf1Δ*, and *pak1-ts rgf1Δ* cells expressing the Rho-probe at permissive (25°C) and restrictive (35.5°C) temperatures.

Stages in cytokinesis progression were determined by measuring the distance between the mitotic SPBs labeled with Sad1-mCherry. We also labeled the actomyosin rings with Rlc1-tdTomato to better discern whether the rings were constricting. Rho1 activation in *pak1+ rgf1+*, *pak1*-*ts rgf1+*, and *pak1+ rgf1Δ* cells at 25°C and at 35.5°C occurred at longer and similar SPB distances (Fig.10B). As previously shown, Rho1 activation occurs in early anaphase in *pak1-ts* cells at the restrictive temperature, (Fig. 8A, B). In agreement with our reasoning, early Rho1 activation in *pak1-ts* mutants at 35.5°C was rescued in the absence of *rgf1* (Fig. 10A, B). These data suggest that early Rho1 activation in the *pak1-ts* mutant is Rgf1-dependent.

**Figure 10.**
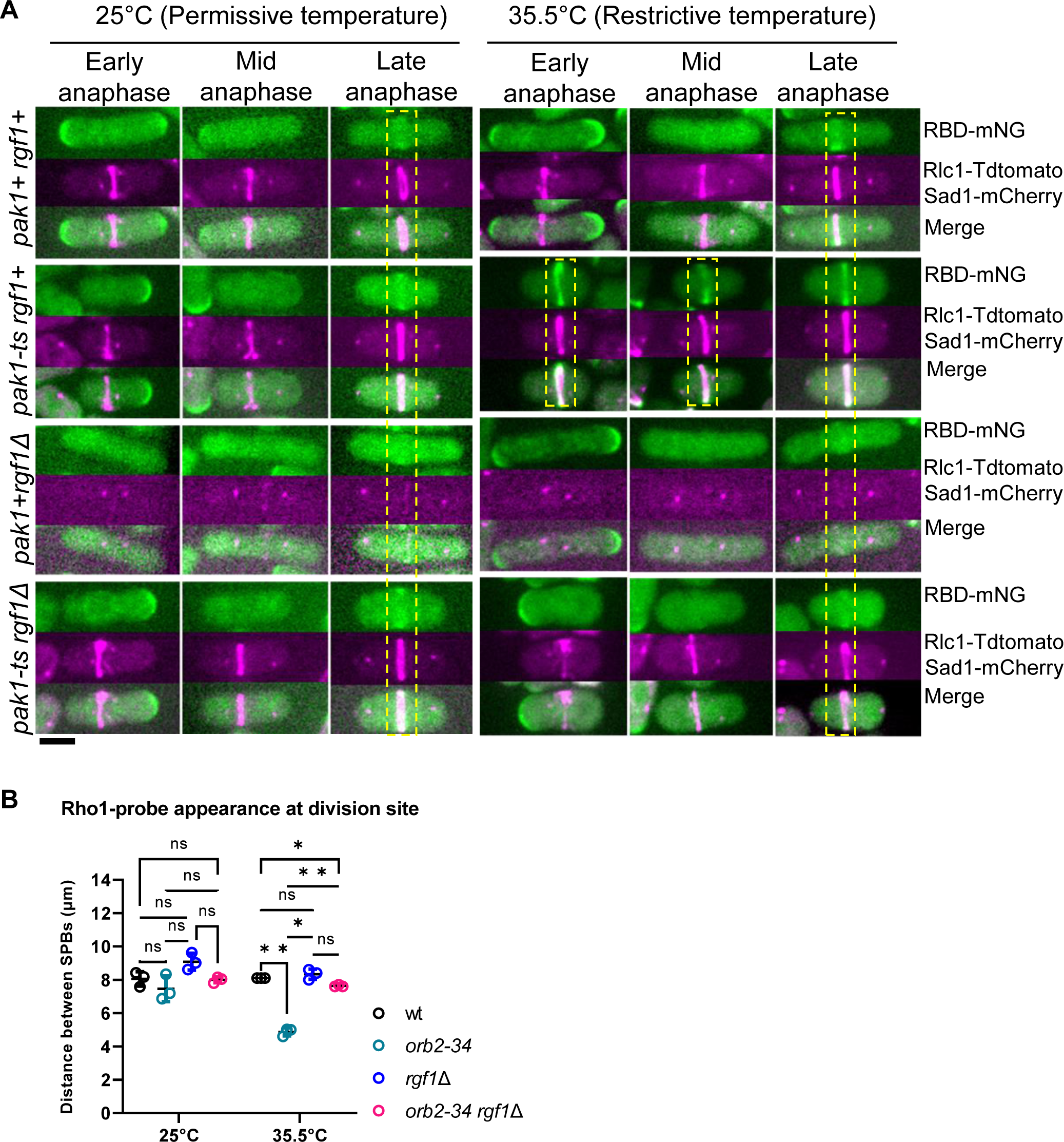
Loss of *rgf1* rescues early Rho1 activation in mutants with disrupted Pak1 function. **A.** Rho1 activation in *rgf1Δ* mutants in *pak1-ts* functional (25°C) and *pak1-ts* hypomorphic (35.5°C) conditions. Yellow boxes highlight stage of cytokinesis in which Rho1 activation is observed in representative cells for each indicated genotype [Scale Bar 5μm]. **B.** Quantification of the distance between the SPBs at which Rho1 activation is observed in all conditions shown. The data points on the graph represent first quartile computations of measurements obtained [N=3 replicate experiments; Statistical significance between strains determined by one-way ANOVA followed by Tukey’s HSD test *p≤0.01,**p≤0.004; n.s *-* not statistically significant; Error bars represent standard deviation].

### Early Rho1 activation leads to cytokinetic defects

What are the implications of early Rho1 activation during cytokinesis? Normally during cytokinesis, the septum is formed in late anaphase, after the actomyosin ring assembles and matures. Further, ring constriction is only initiated when septum deposition starts (Balasubramanian et al., 2004; Proctor et al., 2012). We reported previously that septum ingression and ring constriction begin ∼30minutes after SPB separation, while the recruitment of the primary septum synthesizing enzyme Bgs1 to the division site occurred ∼16minutes after SPB separation (Wei et al., 2016). The Bgs1 enzyme is active and builds the septum only when it binds active Rho1 (Arellano et al., 1996). Thus, the time lag between Bgs1 localization and septum ingression likely occurs due to a delay in Bgs1 activity as a result of Rho1 inhibition in early cytokinesis. Therefore, we postulate that early Rho1 activation in *pak1-ts* mutants will induce early Bgs1 activation, and consequently early septum deposition and ring constriction. In these cells, the time lag between ring assembly and constriction should be alleviated as septum synthesis initiates early. We find that *pak1-ts* mutants displayed a higher number of constricting rings as compared to the controls (Fig. 11A, B, S5A). In addition, these mutants also displayed an increased number of septated cells (Fig. 11C). Taken together, this suggests that ring constriction initiates early in the absence of *pak1* kinase likely due to early septum deposition.

**Figure 11.**
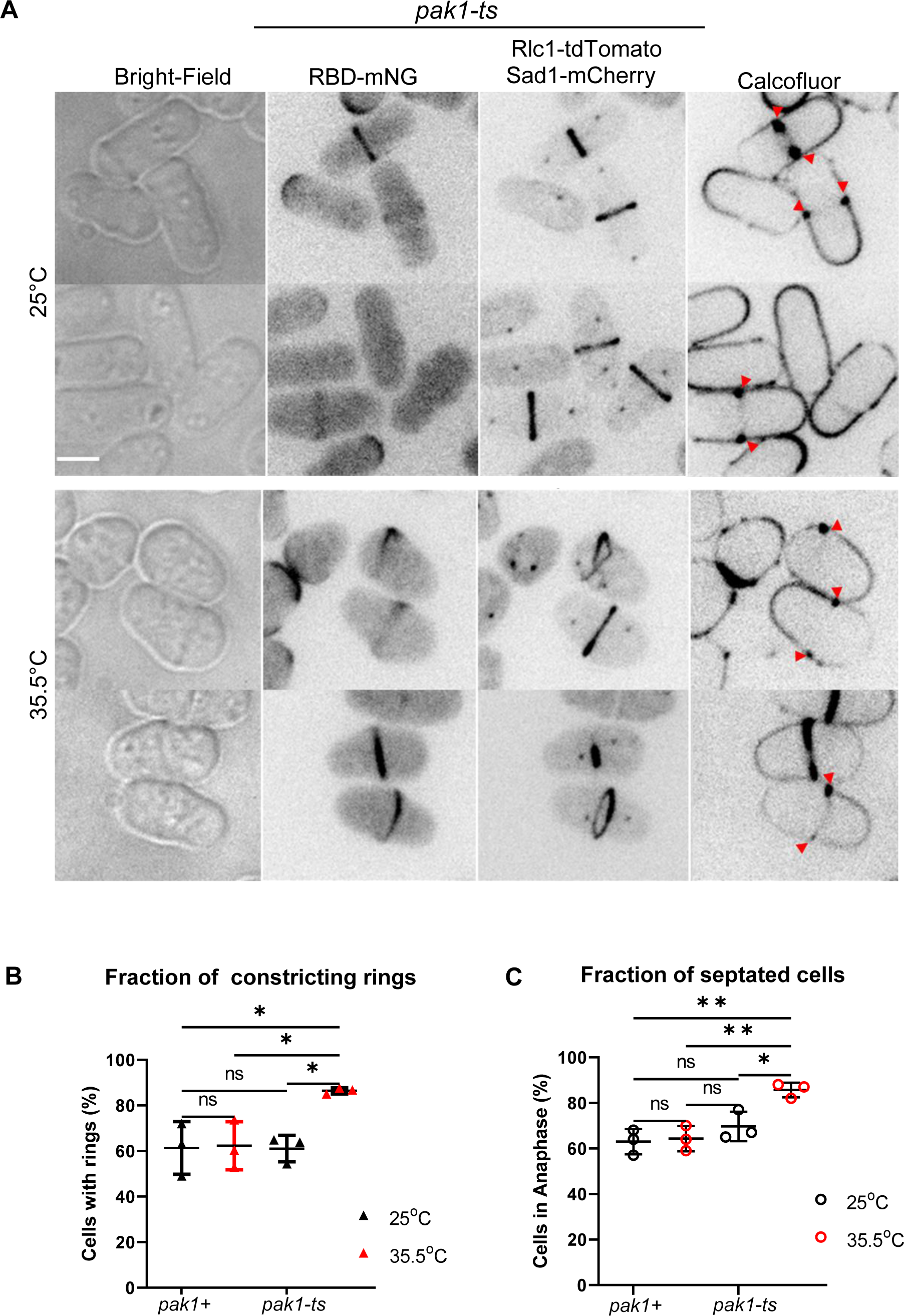
Disruption of Pak1 function results in early septum formation (A-B) **A.** Rho1 activation and septum formation in *pak1-ts* (*orb2-34*) strains grown at permissive and restrictive temperatures. Septum deposition at the division site (red arrowheads) is visualized with calcofluor staining (see methods) [Scale Bar 5μm]. **B.** Quantification of the fraction of constricting rings for cells undergoing cytokinesis in the indicated strains and conditions. [N= 3 representative experiments,*p ≤0.03; n.s *-* not statistically significant; one-way ANOVA, with Tukey’s HSD test; Error bars represent standard deviation]. **C.** Quantification of the fraction of septated cells in the indicated strains and conditions, [N= 3 representative experiments,*p ≤0.025; **p ≤0.005; n.s - not statistically significant; one-way ANOVA, with Tukey’s HSD test; Error bars represent standard deviation].

We quantified the septation index, in *pak1-ts* mutants at 25°C and 35°C. The septation index in an asynchronous wild-type population of cells has been previously reported at ∼18% (Wei et al., 2016). We find that the septation index of *pak1-ts* cells was ∼20%, similar to reported wild-type levels at 25°C. The septation index however doubled to ∼42% when *pak1-ts* mutants were incubated at 35.5°C (Fig. S6. B). Additionally, disruption of *pak1* function caused a small but significant population of dividing cells to become multi-septated (Fig.S6A, C). Our quantifications indicate that ∼3.3% of septated *pak1-ts* mutants at 35.5°C became multi-septated while this was not observed in *pak1-ts* cells at 25°C (Fig. S6A, C). Together these data allow us to propose that active Cdc42 prevents Rho1-activation in early cytokinesis through Pak1-dependent regulation to promote cytokinesis fidelity.

## Discussion

Cytokinesis encompasses events that partition the cytoplasm of dividing cells at the end of the cell cycle. In fission yeast, cytokinesis is accomplished via septum formation and actomyosin ring constriction. The GTPase Rho1 is required for septum formation during cytokinesis and cell wall integrity (Pérez et al., 2018). Previous work indicated that Cdc42 promotes cytokinesis through its roles in septation, membrane trafficking, and concentric furrow formation (Hercyk et al., 2019a; Onwubiko et al., 2019; Onwubiko et al., 2021; Wei et al., 2016). Here, we investigated the relationship between Cdc42 and the essential GTPase Rho1 in the regulation of cytokinesis. We find that while Cdc42 is activated in early anaphase during ring assembly, Rho1 activation occurs in late anaphase, just before initiation of ring constriction. Interestingly, Rho1-specific GEFs Rgf1 and Rgf3 localized very early to the division site, at a time when Rho1 is inactive, suggesting that the GEFs were unable to activate Rho1 in early cytokinesis even though they localize to this site. Our data demonstrate that Cdc42 is responsible for the inhibition of Rho1 activity during early anaphase. In mutants lacking *gef1*, Cdc42 activation at the division site is delayed. In these mutants, Rho1 is activated in early anaphase immediately after the actomyosin ring assembles. We find that constitutively active Cdc42 prevents Rho1 activation at the division site and also at the ends of interphase cells. Together these observations suggest that Cdc42 globally inhibits Rho1 activation both at the division site and the site of cell growth.

The SIN pathway is required for coupling mitosis and cytokinetic events in cells and has been reported to be upstream of Rho1 activation during cytokinesis (Alcaide-Gavilán et al., 2014). Using our active Rho-probe, we showed that the SIN is required for Rho1 activation at the division site and loss of Cdc42 activity cannot bypass this requirement. We also show that the SIN pathway is not required for Rho1 activation at the cell ends. Since Cdc42 inhibits Rho1 both at the division site and the cell ends, it further highlights that the SIN and Cdc42 pathways regulate Rho1 via independent mechanisms.

We find that early Rho1 activation in *gef1Δ* mutants is disrupted only in the absence of the GEF Rgf1. Thus, it is possible that Cdc42 inhibits Rgf1-mediated Rho1 activation during early cytokinesis. We note that Cdc42 activity does not impair or enhance Rgf1 localization to the division site. Thus we posit that Cdc42 only regulates Rgf1’s ability to activate Rho1. While Rgf1 is the primary GEF for Rho1 activation both at the division site and the cell ends, it is not essential (Morrell-Falvey et al., 2005; Mutoh et al., 2005; Tajadura et al., 2004). In contrast, Rgf3 an essential Rho1 GEF only localizes to the division site (Morrell-Falvey et al., 2005; Tajadura et al., 2004). It is not clear why different Rho1 GEFs show different activation patterns at the division site. Our findings suggest that in the absence of active Cdc42, Rgf3 is no longer essential likely due to Rgf1-mediated Rho1 activation.

Reports that regulators of the Rho1 homolog RhoA respond to Pak-mediated regulation have been shown in animal cells (Alberts et al., 2005; DerMardirossian et al., 2004; Tiedje et al., 2008; Zenke et al., 2004). In fission yeast, the p21-activated kinase Pak1 localizes to the division site in early anaphase (Magliozzi et al., 2020). Our observations suggest that Gef1 mediates Rho1 inhibition in early anaphase through a Pak1 kinase-dependent regulation of Rgf1 function. While *pak1* mutants displayed early Rho1 activation, this phenotype was reverted in *pak1 rgf1* double mutants. This supports our hypothesis that Pak1 regulates Rho1 activation via inhibition of Rgf1 function. It is possible that Pak1 phosphorylates Rgf1 or an intermediate protein to block Rho1 activation. Further biochemical analysis will determine the mechanism by which Pak1 blocks Rho1 activation and if this is mediated via Rgf1.

We previously showed that Cdc42 is activated in a Gef1-dependent manner to promote Bgs1 recruitment and for the timely onset of ring constriction (Wei et al., 2016). Thus, while Cdc42 activation promotes the delivery of the first septum synthesizing enzyme, Bgs1, to the division site, it also ensures that this enzyme is not prematurely activated by inhibiting Rho1 (Fig. S7). It is unclear why septum synthesis is tightly regulated during cytokinesis. One potential explanation is to ensure that septum ingression and cell partitioning only occur after completion of nuclear division. Indeed, a recent paper shows that in fission yeast, septum ingression initiates during anaphase B, but at a much slower rate (Garcia Cortes et al., 2018). The rate of septum ingression increases only after the completion of anaphase B. Inhibition of Rho1 activation likely allows this careful coordination of mitosis and septum ingression. In *gef1Δ* cells, the delay in Bgs1 recruitment in the absence of active Cdc42, even as Rho1 is activated early, ensures a delay in septum ingression. In contrast, the *pak1* mutants appear to initiate septum ingression early, likely due to proper Bgs1 delivery and premature Rho1 activation. Our findings may explain the previously reported observation that *pak1* kinase defective mutants prematurely initiate ring constriction (Loo and Balasubramanian, 2008). Thus, active Cdc42 at the division site in anaphase enforces spatiotemporal regulation of β-glucan septum synthesis. Cdc42 therefore functions as a cellular quality-control at the division site to facilitate time-dependent organization of cytokinetic events.

In animal cells RhoA activation is essential for the formation of an actomyosin ring (Basant and Glotzer, 2018). RhoA is activated at the division site and this leads to actomyosin ring formation at that site (Wagner and Glotzer, 2016). However, in fission yeast Rho1 is activated at the division site only after the ring is fully assembled and ready to constrict. This activation pattern does not support a role for Rho1 in ring assembly but supports a role in septum formation. Our work further provides new details of Cdc42 and Rho1 crosstalk during fission yeast cytokinesis, where these GTPases localize in concentric zones to regulate essential cytokinetic steps. While we show that Cdc42 inhibits Rho1 activation in early anaphase, this inhibition is finally removed in late anaphase to enable septum formation. Concentric zones of active Cdc42 and RhoA have also been reported to drive wound healing in *Xenopus* oocytes (Benink and Bement, 2005). Crosstalk between Cdc42 and RhoA maintains these zones of GTPase activity and constriction dynamics. Indeed, it was shown that dominant-negative Cdc42 eliminates RhoA activation during constriction of the actomyosin array in wound healing, while constitutively active Cdc42 broadens the RhoA activity zone (Benink and Bement, 2005). The specific zones of activity for Rho-GTPases may be required for regulating distinct cellular processes and may be determined via regulation of their GEFs and GAPs to maintain GTPase distinct zones. It will be interesting to assess whether Rho1-dependent effectors regulate active Cdc42 zones during cytokinesis.

During growth in fission yeast, Cdc42 is restricted to growing tips primarily via the activity of its GAP Rga4 (Das et al., 2007; Rich-Robinson et al., 2021). At the onset of ring constriction during cytokinesis, Cdc42 GAPs also localize to the division site (Campbell et al., 2021; Rich-Robinson et al., 2021). These GAPs may establish distinct zones that are devoid of active Cdc42 where Rho1 activation can occur to finally allow septum formation. In budding yeast, antagonism between Cdc42 and Rho1 regulates secondary septum formation, and inactivation of Cdc42 is required for proper cytokinesis completion (Atkins et al., 2013; Onishi et al., 2013). In-depth biochemical analysis and mathematical models that probe toggles between GTPase regulators in concentric zones will provide insights into the mechanisms of crosstalk between these Rho GTPases.

## Materials and Methods

### Strains and cell culture

Strains used in this study are listed in Table S1. The fission yeast strains used in this study are isogenic to PN972. Unless mentioned, cells were cultured in yeast extract (YES) medium and grown exponentially at 25°C. All genetic manipulations of strains were carried out using standard techniques (Moreno et al., 1991). Cells were grown exponentially for at least three rounds of eight generations before experiments were performed.

### Microscopy

Image acquisition was performed at room temperature (23–25°C) on a spinning disk confocal system that uses a Nikon Eclipse inverted microscope with a 100× 1.49NA objective, a CSU-22 spinning disk system, and a Photometrics EM-CCD camera from Visitech International. Images were acquired using Metamorph (Molecular Devices) and analyzed using ImageJ/FIJI Bio-Formats plugins (National Institutes of Health). For still images, cells were mounted on glass slides with a #1.5 coverslip (Fisher Scientific, Waltham, MA) and imaged right away. All Z-series images were acquired with a depth interval of 0.4 µm for a total of 6.2 µm. In time-lapse image acquisition, cells were placed in a 3.5-mm glass-bottom culture dish and covered with YES medium with 0.6% agar. Ascorbic acid (100 µM vitamin C) was added to the cell culture to minimize fluorescence toxicity, as previously reported (Wei et al., 2017). All images analyzed for mean intensity were acquired with Z-series and sum-projected unless noted otherwise. Statistical analysis was performed using one-way ANOVA, followed by Tukey’s honestly significant difference (HSD) posthoc test or Student’s t-test where appropriate. Comparisons between experimental groups were considered significant when P≤0.05.

### Designing the active Rho-probe

The Rho-probe was designed using the Rho-Binding Domain of the yeast protein kinase C (*pkc2*) used in (Kono et al., 2012). We modified the expression of RBD by using the *pkc2* promoter region from -464 to -1 from the start codon to drive its expression. We stitched three fragments: pkc2 promoter sequence, the RBD, and mNG-pjk148 or tdTomato-pjk210 using restriction sites in our primer design. The C-terminal sequence of the RBD contains a glycine linker with either mNeonGreen or tdTomato in a Pjk148 or Pjk210 vector. Cells were transformed via lithium acetate transformation (Okazaki et al., 1990) to integrate constructs into the leu, or *ura* locus of PN975/YMD493.

Primer sequences used for the *pkc2* promoter region were as follows-

**Fwd:** 5’-AAG CTT GAT ATC GAA TTC CTG CAG CCC GGG AAT GAA CTG TTC TAT TAA TTG GTC-3’

**Rev:** 5’-GTG AAA TCA TTA CTT TAA GCC TAA TCC-3’

### Cytoskeleton disruptions

To block Arp2/3 complex-dependent branched actin assembly, cells were treated with 100 µM CK666 (Sigma-Aldrich, SML006-5MG) in dimethyl sulfoxide (DMSO, Sigma-Aldrich, D8418-250ML). To disassemble all F-actin, cells were treated with 100µM Latrunculin A (LatA-EMD Millipore) dissolved in dimethyl sulfoxide (DMSO) in YE media for 30minutes before imaging. To depolymerize microtubules, cells were treated with 25µg/ml MBC dissolved in DMSO and incubated for 45 minutes before imaging. For all these experiments, control cells were treated with 0.1% DMSO in YES media.

### Expressing constitutively active Cdc42

The *cdc42G12V* fragment is cloned into the *pJK148* vector under the thiamine-repressible promoter nmt41 and integrated into the genome of *gef1+* and *gef1Δ* cells, as previously described in Wei et al. (2016). Cells were initially grown in EMM (Edinburgh minimal medium) with 15 µM thiamine. Partial induction of cdc42G12V expression was performed by harvesting strains via low-speed centrifugation, rinsing 4 times with deionized water, and then grown in EMM with 0.05µM thiamine for 34hours before imaging at 25°C. The experimental controls were *gef1+* and *gef1Δ* cells transformed with the empty pJK148 vector.

### Quantification of fluorescence intensity

Fluorescence intensity was measured in images via ImageJ software. All images were sum-projected and mean intensities were reported. A box was positioned to measure the signal at the division site. The cytoplasm of the cell with little to no signal was used for background subtraction. Mean intensity measurements were collected after background subtractions.

### Estimation of timing of cytokinetic progression

To determine the timing of cytokinetic progression, we measure the spindle pole body distance in cells undergoing mitosis. Mitotic progression as measured via spindle pole body distance functions as an internal clock for cytokinetic events. The timepoint at which the two spindle poles can be first distinguished is zero time. Z-series images of cells expressing the spindle pole body protein Sad1-mCherry and the cytokinetic ring marker Rlc1-tdTomato were acquired for cells in all conditions measured. The Line tool in ImageJ was used to measure the distance in microns (µm) between the spindle pole bodies (SPBs) in cells. Z-series helped visualize spindles that were on different focal planes. Rlc1-tdTomato signal was also helpful for the clarification of constricting rings. Since SPB measurements were obtained in an asynchronous population of cells, we only plotted the 25^th^ percentile of our SPB data measurements which represents the smallest SPB measurements for each strain in each data set. All experiments were performed in triplicate.

### SIN inactivation via *sid2-250*-ts

*sid2-250* temperature-sensitive cells were cultured in YE media until healthy. Cells were grown to an OD of 0.2 at 25°C, and culture split into two halves, for each strain. One half was kept at the permissive temperature of 25°C for 4 hours prior and then imaged. The other half was shifted to the restrictive temperature of 36°C for 4 hours to inactivate Sid2 and then imaged as previously performed (Feoktistova et al., 2012).

### *rgf3*-repression

To repress *rgf3,* nmt-81-*rgf3* cells *(VT88)* were grown in supplemented EMM (Edinburgh minimal medium). Weak suppression of *rgf3* was achieved by growing nmt-81-*rgf3* cells in EMM +Sup media with 15 µM thiamine for 6 generations as previously performed (Tajadura et al., 2004). For controls, cells were grown in EMM supplemented media with no thiamine for the same amount of time.

### Spot Growth assay

Cells were grown to OD 0.5 (1x10^7^cells/μL), in EMM media supplemented with Adenine, Leucine, Histidine, and Uridine, (A, L, U, H) at 25°C. Serial dilutions were set up from 10^4^ to 10^1^ cells, which were spotted for each strain on both EMM-ALUH + 0μM thiamine, and EMM-ALUH + 15μM thiamine media plates and incubated at 25°C. Cell growth was assessed after 7 days.

### Disruption of Pak1 kinase function via temperature-sensitive allele *orb2-34*

Pak1 kinase was inactivated via the *orb2-34* (*pak1-ts*) temperature-sensitive mutation. Cells were grown till healthy in YE media. On the day of the experiment, cells were cultured to an OD of 0.2. A subset of these cells were incubated at the permissive temperature of 25°C, while the other set was incubated at the restrictive temperature of 35.5°C for 4hours. Cells were imaged after 4 hours.

### pak1 overexpression pak1+, (pak1OE), and pak1 repression (pak1-)

In experiments with *nmt1-3HA-pak1* (MBY), cells were grown in YE media for at least 3 generations. Cells were washed thoroughly 4 times with thiamine-free EMM media, switching to a new tube during the fourth wash. Washed cells were divided into two groups: For repression of *nmt1-3HA-pak1*, washed cells were transferred to EMM media with 15uM thiamine and grown for 48hours then imaged. For *pak1OE*, washed cells were transferred to EMM media containing no thiamine and grown for 48 hours, and imaged immediately.

### Calcofluor staining

To stain the septum and cell wall, live cells were stained in YE liquid with 50 μg/ml Calcofluor White M2R (Sigma-Aldrich, St. Louis, MO) at room temperature and imaged.

## ACKNOWLEDGEMENTS

We thank Yolanda Sanchez, Kathy Gould, Mohan Balasubramanian, and Pilar Perez for providing strains.

