## Supplemental Material for "Cdc42 prevents precocious Rho1 activation during cytokinesis in a Pak1-dependent manner"

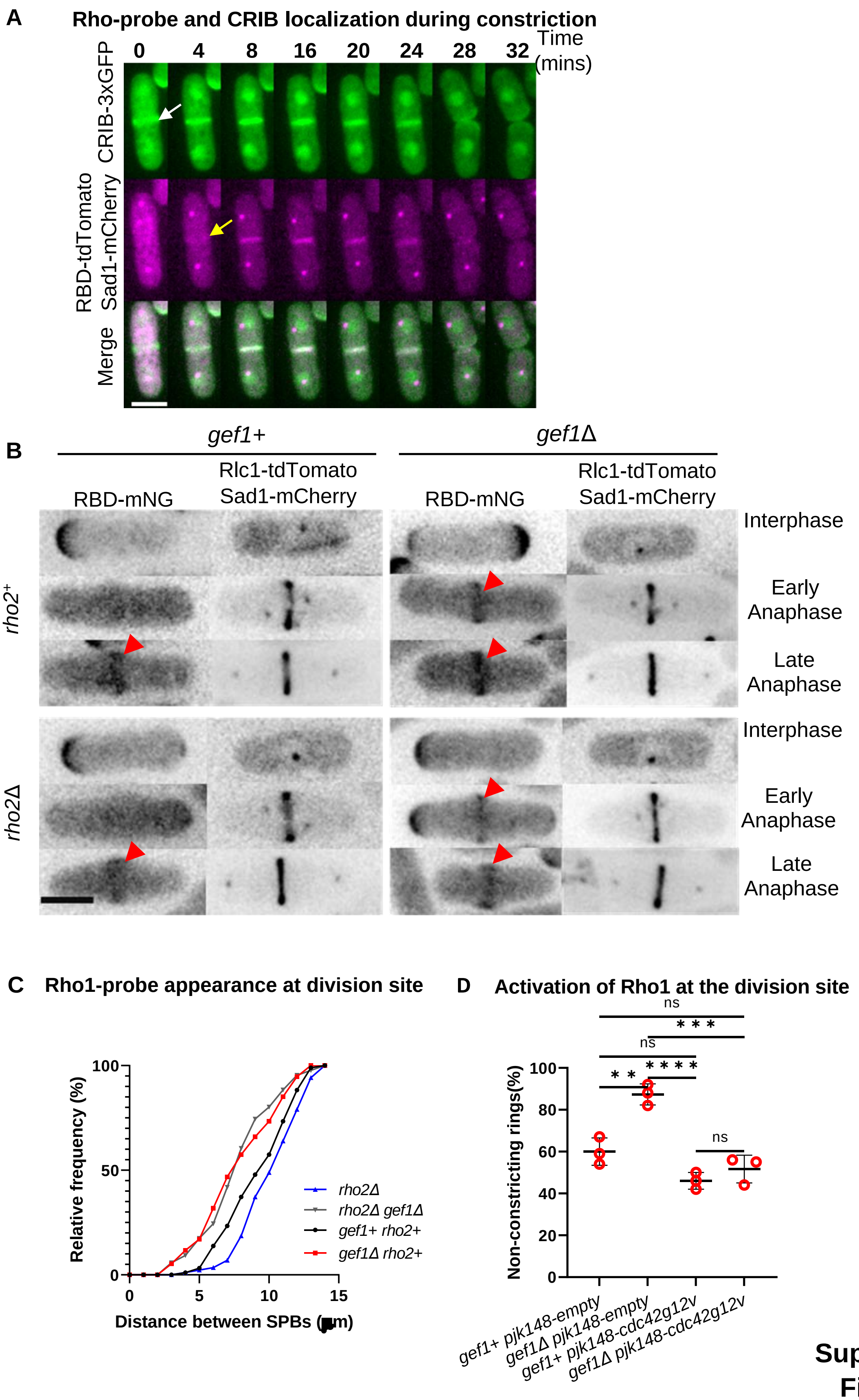


**Supplement Figure 1**

**Figure S1. Active Rho-probe detects Rho1 activation in cells**

**A.** Time-lapse montage of a representative cell showing Rho1 (yellow arrow) and Cdc42 activation (white arrow) at the division site during ring constriction [Scale Bar 5μm]. **B.** Sum projection of z-stack images showing that the Rho-probe detects active Rho1 in *gef1+ rho2+* and *gef1Δ rho2Δ* strains. Red arrowheads point to active Rho1 localization in dividing cells [Scale Bar 5μm]. **C.** Outcome plot shows the SPB distances at which active Rho1 is observed at the division site in the mentioned strains, n=103 cells per indicated strains. **D.** Quantification of the percentage of non-constricting rings in strains as indicated. [N=3 experiments; Statistical significance between strains determined by one-way ANOVA followed by Tukey's HSD test, $^{**}p \leq 0.0016$ $^{***}p \leq 0.0003$, $^{****}p \leq 0.0001$; n.s - not statistically significant; Error bars represent standard deviation].

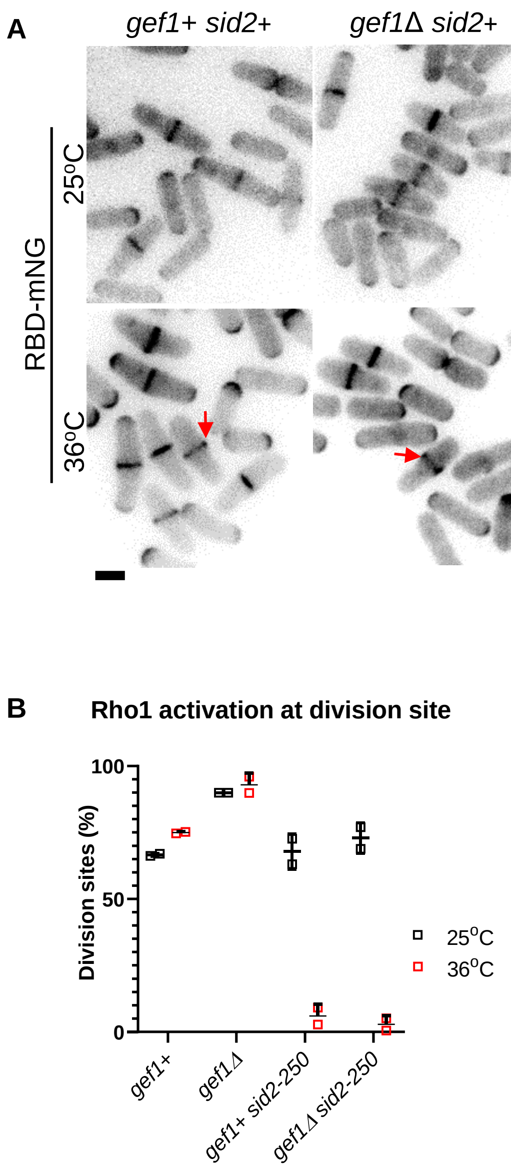


**Figure S2. Proper function of the SIN pathway is required for Rho1 activation at the division site**

**A.** Rho1 activation during cytokinesis in *sid2-250* strains incubated for 4hours at the permissive temperature (25°C) and restrictive temperature (35.5°C) [Scale Bar 5μm]. Red arrows point to division sites. **B.** Quantification of the fraction of division sites with Rho1-probe localization [N= 2 experiments, (≥100 division sites analyzed per strain for each experiment); Error bars represent standard deviation].

**Supplement Figure 2**

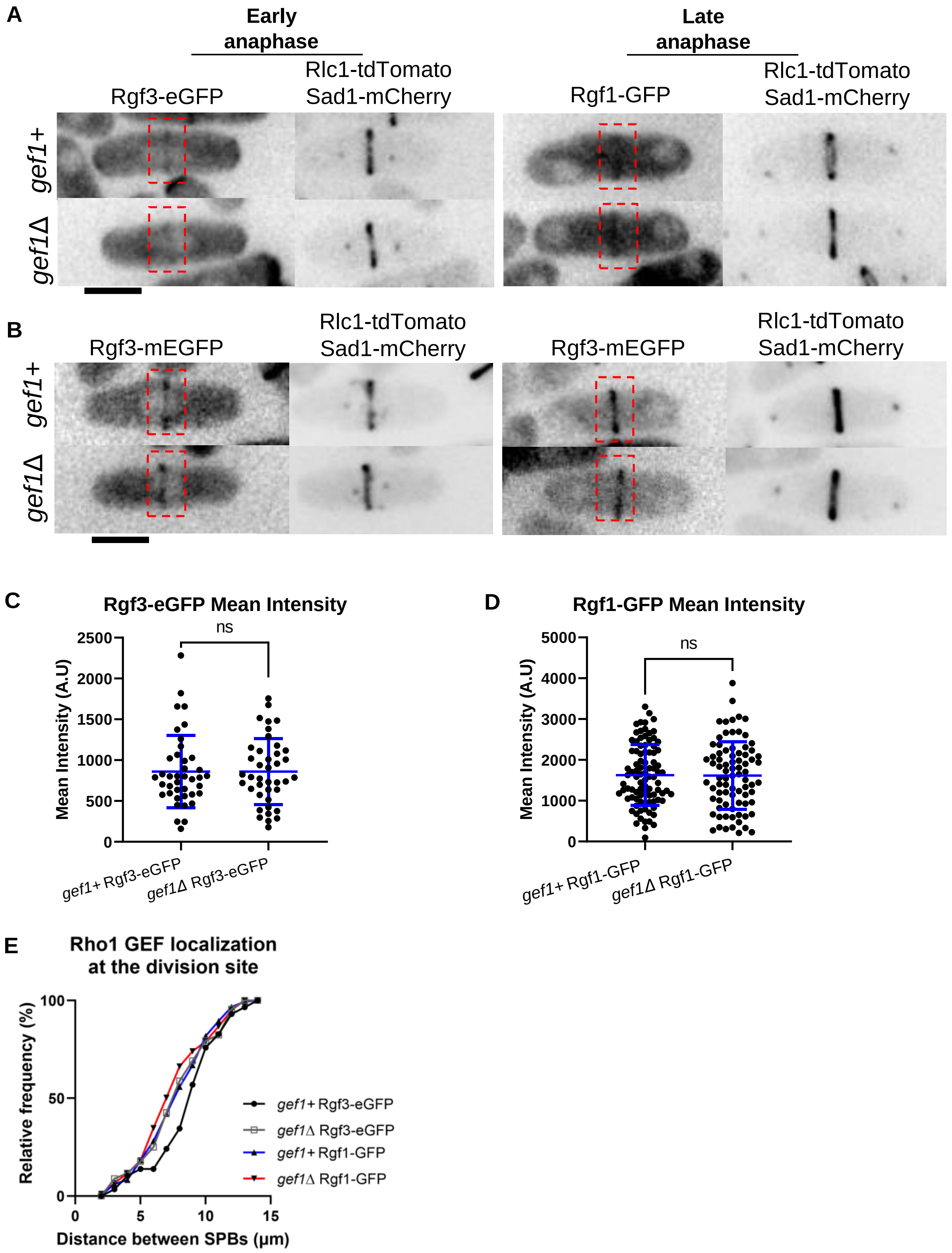


**Supplement Figure 3**

**Figure S3. Localization of Rho1 GEFs is similar in *gef1*+ and *gef1Δ* cells**

Sum projections of cells showing localization of Rgf3-eGFP **(A),** and Rgf1-GFP **(B)** to the division site during early and late cytokinesis (red boxes) in the *gef1*+ and *gef1Δ* cells. Quantification of mean fluorescent intensities of Rgf3-mE-GFP **(C),** and Rgf1-GFP **(D),** at the division site of indicated strains, [n ≥80 division sites analyzed per strain; Statistical significance between strains determined by Mann-Whitney test, Rgf3 p=0.774, Rgf1 p=0.936, n.s - not statistically significant; Error bars represent standard deviation]. **E.** Outcome plot shows the distance between the SPBs at which Rgf3 and Rgf1 localization are observed at the division site, n≤190 cells per indicated strains.

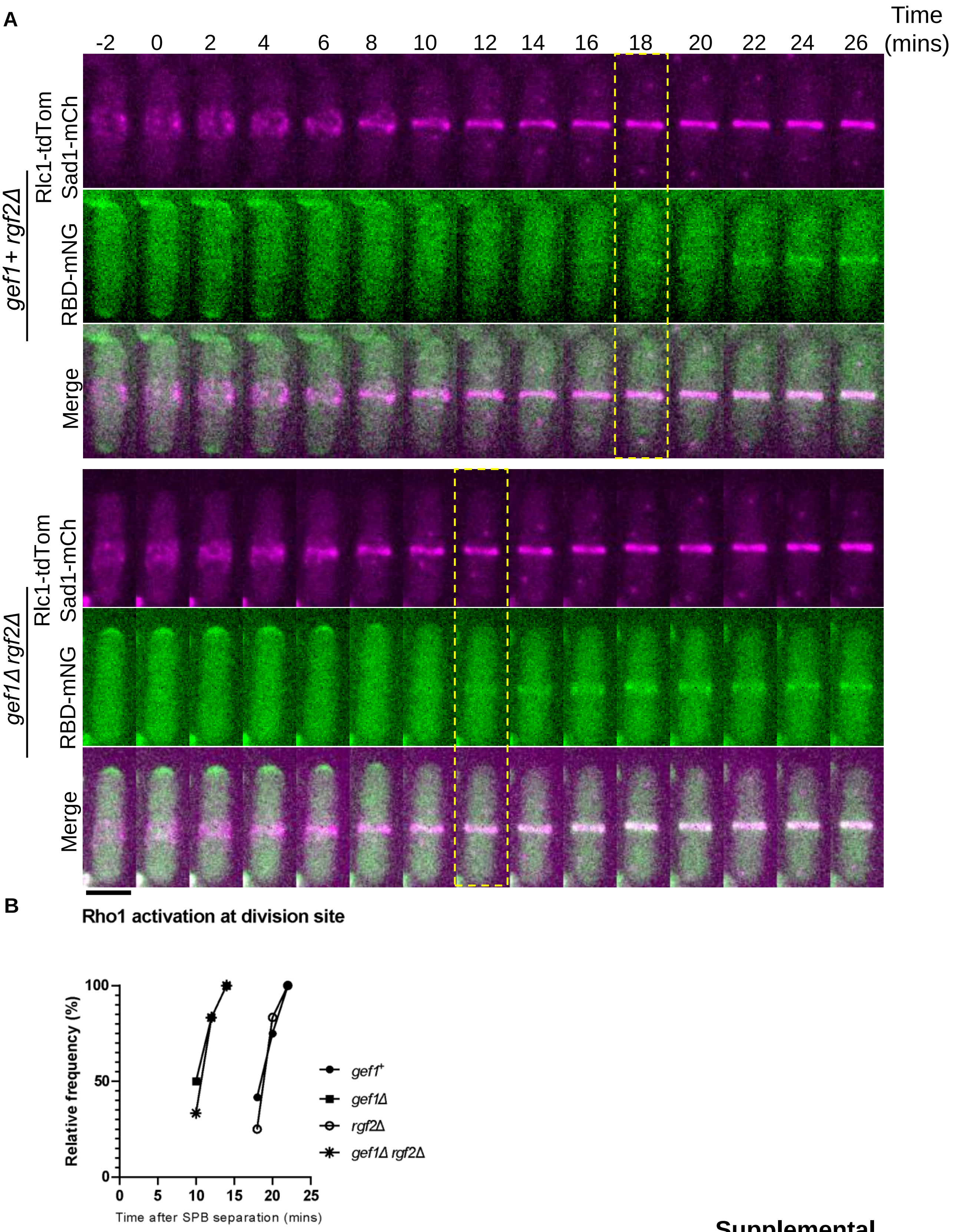


**Supplemental Figure 4**

**Figure S4. Loss of *rgf2* does not rescue Rho1 activation in *gef1* mutants**

**A.** Time-lapse of representative *gef1+ rgf2+*, and *gef1Δ rgf2Δ* cells shows the time of Rho1 activation at the division site during cytokinesis (yellow box) [Scale Bar 5µm Time=0 marks the time of SPB separation, and onset of cytokinetic events. **B.** Outcome plot shows the frequency of Rho1 activation over time at the division site during cytokinesis [n=12 cells per strain]. **C.** Quantification of the timing of Rho1 activation during cytokinesis from movies of the listed strains, [n=12 cells per strain; Statistical significance between strains determined by one-way ANOVA followed by Tukey's HSD test ****$p \leq 0.0001$; n.s - not statistically significant; Error bars represent standard deviation].

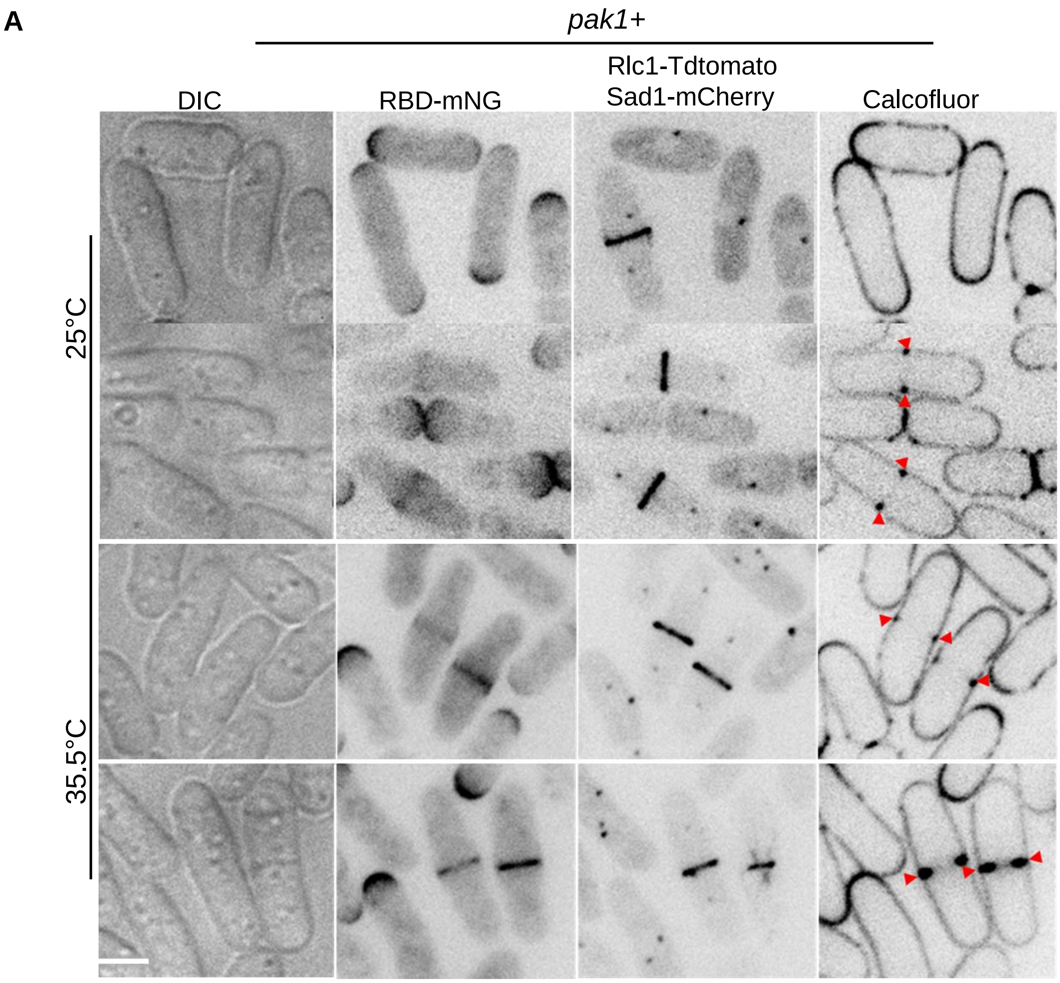


**Figure S5. Hypomorphic *pak1* mutant results in early Rho1 activation in cells during cytokinesis**

**A.** Rho1 activation and septum formation in *pak1+* (*orb2+)* strains grown at permissive (25ºC) and restrictive temperatures (35.5ºC). Septum deposition at the division site (red arrowheads) is visualized with calcofluor staining (see methods) [Scale Bar 5µm].

**Supplemental Figure 5**

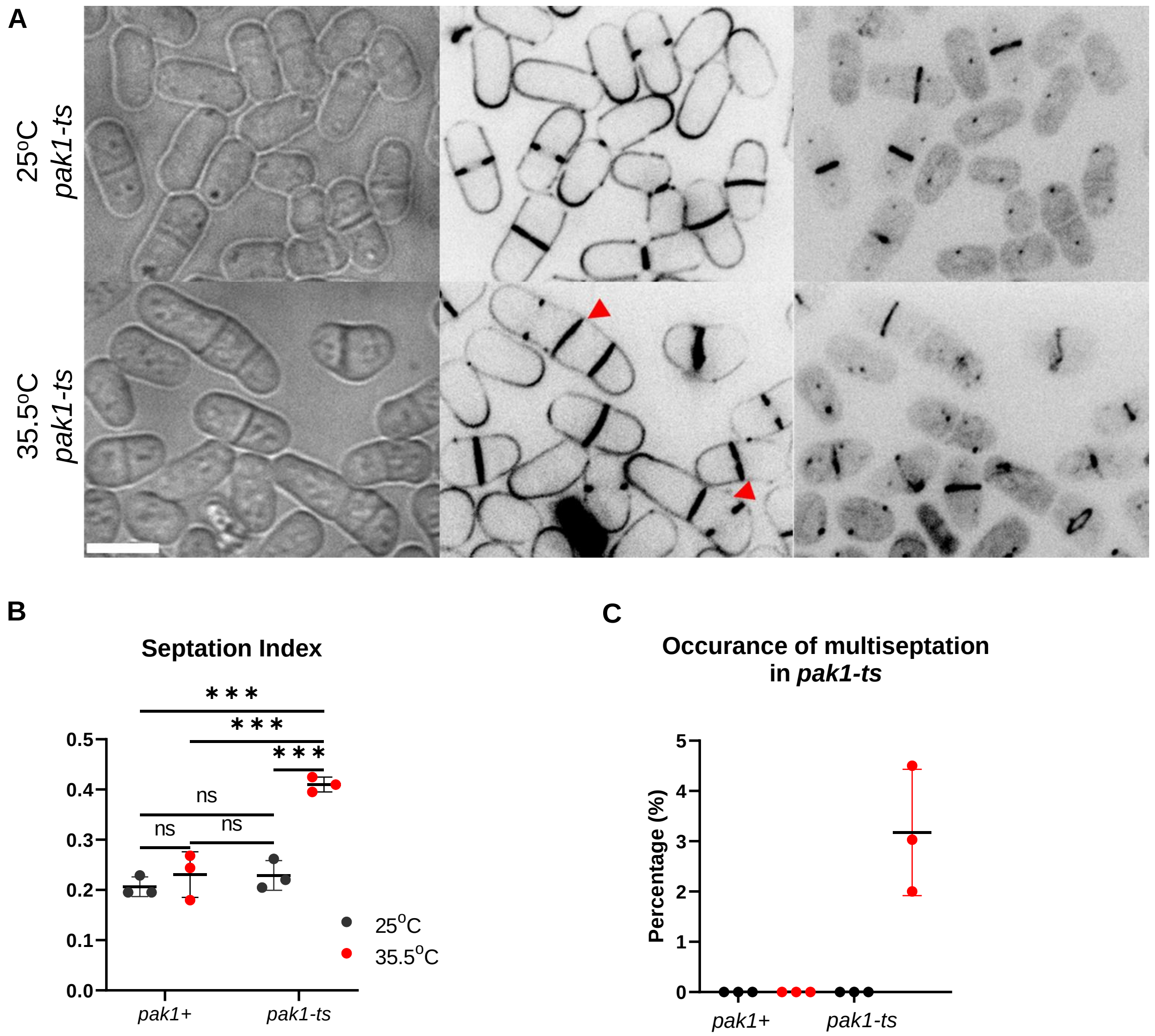


**Figure S6. Cells with early Rho1 activation exhibit early septum formation and ring constriction, and display cytokinetic defects**

**A.** *pak1-ts* mutants at the restrictive temperature (35.5°C) become multi-septated, red arrowheads compared to control experimental conditions (25°C) [Scale Bar, 5µm]. **B.** Quantification of the septation index in an asynchronous population of *pak1+* and *pak1-ts* strains at 25°C and 35.5° C. [N=3 experiments, Statistical significance between strains determined by Student's t-test, ***p≤0.0005; Error bars represent standard deviation]. **C.** Quantification of the percentage of multi-septated cells in the indicated strains when cells are incubated at 35.5°C, [N=3 replicate experiments; Error bars indicate standard deviation].

**Supplemental Figure 6**

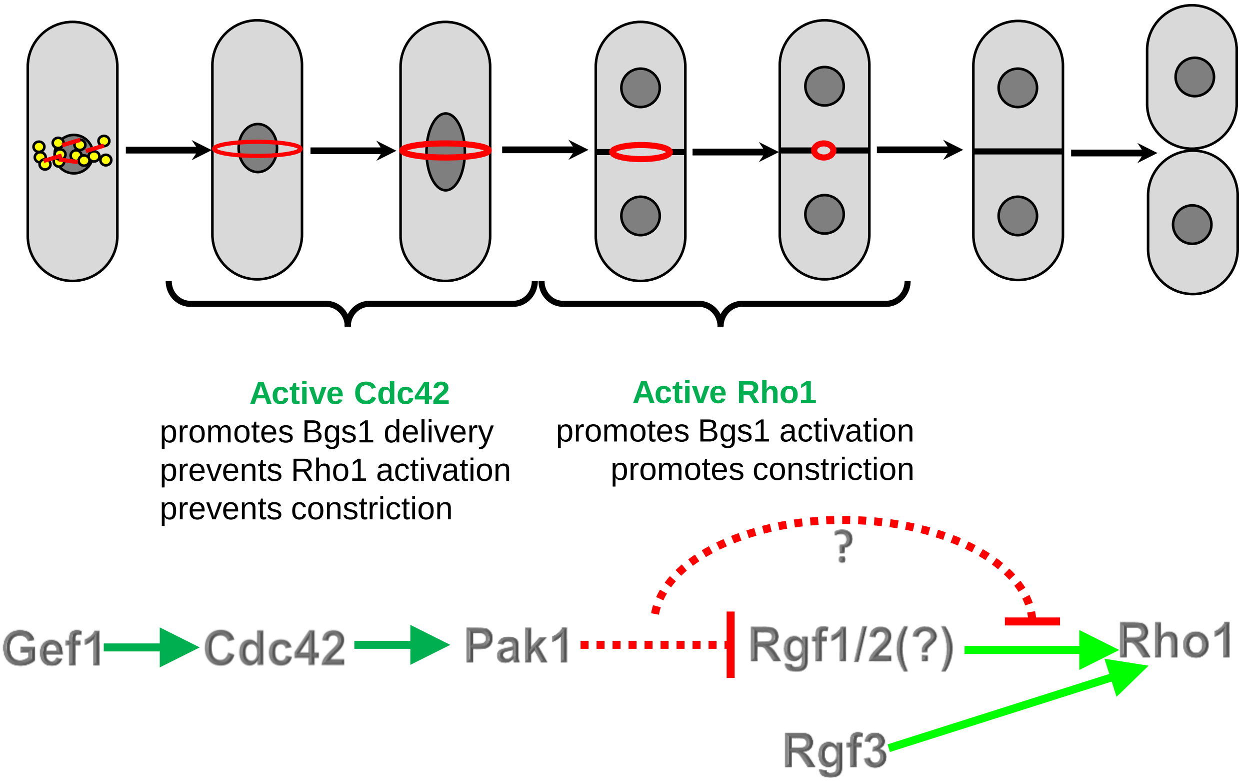


**Figure S7.**

Schematic describing how Cdc42 prevents early Rho1 activation and thus promotes proper cytokinesis fidelity. Gef1 activates Cdc42 which in turn activates the Pak1 kinase. The Pak1 kinase inhibits Rho1 activation.

**Supplemental Figure 7**

**Strain list**

**Table S1**

| Strain | Genotype | Origin |
|---|---|---|
| PN975 YMD493 | *h+ ura4-D18 leu1-32 ade6-704* | P. Nurse |
| YMD527 | *Rlc1-tdTomato-NATr Sad1-mCherry: kanMx ade6-M21X leu1-32 his7+ ura4-D18* | This study |
| YMD1062 | *leu2: pck2:RBD-Neon Green leu+ Rlc1-tdTomato-NATr Sad1-mCherry: kanMx* | This study |
| YMD1099 | *gef1Δ::ura+ leu: pck2:RBD-Neon Green leu+ Rlc1-tdTomato-NATr Sad1-mCherry: kanMx* | This study |
| YMD1394 | *orb2-34 (pak1-ts) leu:pck2:RBD-Neon Green leu+ Rlc1-tdTomato-NATr Sad1-mCherry: kanMx* | This study |
| YMD1706 | *Rgf1-GFP:KanMx Rlc1-tdTomato: NATr Sad1-mCherry:KanMX* | This Study |
| YMD1697 | *gef1Δ::ura+ rgf2Δ::KanMx leu2: pck2:RBD-mNeonGreen:leu+ Rlc1-tdTomato-NATr Sad1-mCherry: KanMx* | This Study |
| YMD1729 | *rgf2Δ::Kan leu: pck2:RBD-mNeonGreen:leu+ Rlc1-tdTomato-NATr Sad1-mCherry:KanMx* | This Study |
| YMD1699 | *gef1Δ::ura+ rgf1Δ::KanMx leu2: pck2:RBD-mNeonGreen:leu+ Rlc1-tdTomato-NATr Sad1-mCherry: KanMx* | This Study |
| YS733 | *h- rgf1Δ::KanMx* | Gift from Y.Sanchez |
| YS2147 | *h- rgf2Δ::KanMx* | Gift from Y.Sanchez |
| PPG0378 | *h- rga5Δ::ura+ leu1-32* | Gift from P.Perez |
| YMD1728 | *rgf1Δ::KanMx leu: pck2:RBD-mNeonGreen:leu+ Rlc1-tdTomato-NATr Sad1-mCherry KanMx* | This Study |
| YMD1786 | *p3-nmt1-3xHA shk1:G418 leu2: pck2:RBD-mNeonGreen:leu+ Rlc1-tdTomato-NATr Sad1-mCherry: KanMx* | This Study |
| YMD1801 | *pak2Δ::Kan leu: pck2:RBD-mNeonGreen:leu+ Rlc1-tdTomato-NATr Sad1-mCherry: KanMx* | This Study |
| YMD1775 | *gef1Δ::ura+ p3-nmt1-3xHA shk1:G418 leu: pck2:RBD-mNeonGreen:leu+ Rlc1-tdTomato-NATr Sad1-mCherry: kanMx* | This Study |
| VT88 | *rgf3Δ(nmt81-rgf3+)* | Gift from Y.Sanchez |
| YMD1823 | *rgf3Δ(nmt81-rgf3+) leu:pck2:RBD-mNeonGreen:leu+ Rlc1-tdTomato-NATr Sad1-mCherry: kanMx* | This Study |
| YMD1717 | *gef1Δ::ura+ rgf3Δ(nmt81-rgf3+) leu:pck2:RBD-mNeonGreen:leu+ Rlc1-tdTomato-NATr Sad1-mCherry: kanMx* | This Study |
| YMD1714 | *nmt41-cdc42g12v:leu+ RBD-tdTomato:ura+ Rlc1-GFP:Kan Sad1-mCherry:KanMx* | This Study |
| YMD1715 | *gef1Δ::ura+ Rgf1-GFP:Kan Rlc1-tdTomato: NATr Sad1-mCherry:KanMX* | This Study |

| | | |
|---|---|---|
| YMD1827 | *rho2Δ::ura+ leu:pck2:RBD-mNeonGreen:leu+ Rlc1-tdTomato-NATr Sad1-mCherry: KanMx* | This Study |
| YMD1826 | *gef1Δ::ura+ rho2Δ::ura+ leu:pck2:RBD-mNeonGreen:leu+ Rlc1-tdTomato-NATr Sad1-mCherry: kanMx* | This Study |
| YMD1628 | *Mob1-mEGFP: KanMx Rlc1-tdTomato: NATr* | This Study |
| YMD1629 | *Sid2-mEGFP: KanMx Rlc1-tdTomato: NATr* | This Study |
| YMD1636 | *gef1Δ:ura+ Mob1-mEGFP:KanMx Rlc1-tdTomato:NATr* | This Study |
| YMD1635 | *gef1Δ:ura+ Sid2-GFP:KanMx Rlc1-tdTomato:NATr* | This Study |
| YMD1632 | *h+ gef1Δ:ura+ nmt41-pjk148-empty: leu+ gef1Δ:ura+ RBD-tdTomato:ura+ Rlc1-GFP:KanMx Sad1-mCherry:KanMx* | This Study |
| YMD1602 | *nmt41-pjk148-empty: leu+ gef1Δ:ura+ RBD-tdTomato:ura+ Rlc1-GFP:KanMx Sad1-mCherry:KanMx* | This Study |
| YMD1616 | *gef1Δ:ura+ nmt41-cdc42g12v: leu+ gef1Δ::ura+ RBD-tdTomato:ura+ Rlc1-GFP:KanMx Sad1-mCherry:KanMx* | This Study |
| YMD1045 | *leu2:pck2:RBD-mNeonGreen:leu+* | This Study |
| YMD1493 | *sid2-250 leu2:pck2:RBD-mNeonGreen:leu+ Rlc1-tdTomato-NATr Sad1-mCherry: KanMx* | This Study |
| YMD1491 | *gef1Δ::ura+ sid2-250 leu:pck2:RBD-mNeonGreen:leu+ Rlc1-tdTomato-NATr Sad1-mCherry: KanMx* | This Study |
| YMD1119 | *Rgf3-mEGFP:leu+ Rlc1-tdTomato: NATr Sad1mCherry:KanMX* | This Study |
| YMD1121 | *gef1Δ::ura+ Rgf3-mEGFP:Kan Rlc1-tdTomato: NATr Sad1-mCherry:KanMX* | This Study |
| YMD764 (MBY3451) | *h- nmt1-3xHA-pak1* | Loo et al., 2008 |
| YMD1708 | *rga5Δ leu:pck2:RBD-mNeonGreen:leu+ Rlc1-tdTomato-NATr Sad1-mCherry: KanMx* | This Study |
| YMD1652 | *gef1Δ::ura+ rga5Δ:: ura+ leu:pck2:RBD-mNeonGreen:leu+ Rlc1-tdTomato-NATr Sad1-mCherry: KanMx* | This Study |
